# Chemogenetic profiling of ubiquitin-like modifier pathways identifies NFATC2IP as a mediator of SUMO-dependent genome integrity

**DOI:** 10.1101/2023.06.30.547196

**Authors:** Tiffany Cho, Yichao Zhao, Michele Olivieri, Lisa Hoeg, Dheva Setiaputra, Daniel Durocher

## Abstract

The post-translational modification of proteins by ubiquitin and ubiquitin-like polypeptides controls multiple cellular processes including the abundance of a large fraction of the proteome. We applied genome-scale CRISPR/Cas9 screens to elucidate the genetic architecture of the response to inhibition of ubiquitin, NEDD8 and SUMO conjugation pathways as well as inhibition of the p97/VCP segregase. This effort identified 395 genes whose disruption alters the fitness of human cells when faced with perturbations in these pathways. We validated that the TMED2 and TMED10 proteins, which are localized to the secretory pathway, promote resistance to p97/VCP inhibition and also characterized NFATC2IP, an evolutionarily conserved protein harboring SUMO-like domains as a major player in promoting genomic integrity when SUMOylation is inhibited. We propose that NFATC2IP acts in interphase cells to promote the SUMO-dependent E3 ligase activity of the SMC5/SMC6 complex, which is critical for SUMO-dependent genome integrity.

## Introduction

The covalent attachment of ubiquitin and ubiquitin-like (Ubl) polypeptides to target proteins and other macromolecules modulate myriad cellular processes (Hershko and Ciechanover 1998; Dikic and Schulman 2022). By far the best characterized post-translational modification in this family is ubiquitin conjugation with the subsequent formation of ubiquitin chains that act as signals for degradation by the 26S proteasome (Finley 2009). Ubiquitin-dependent degradation controls the abundance of many signaling and cell cycle proteins and controls processes such as the termination of DNA replication and ribosome quality control (Hoeller and Dikic 2009; Dewar and Walter 2017; Joazeiro 2017). Ubiquitin also acts as a signaling or organizing molecule in processes such as DNA repair and receptor trafficking where non-degradative ubiquitin chain topologies are used to coordinate protein-protein interactions (Jackson and Durocher 2013; Foot et al. 2017).

Ubiquitin conjugation requires a multi-enzymatic cascade that is initiated by an E1 activating enzyme that uses ATP to form a high energy ubiquitin∼E1 thioester intermediate. Ubiquitin is then transferred to an E2 conjugating enzyme, which is then poised to conjugate the ubiquitin moiety to a substrate, usually by forming an isopeptide bond with the ε-amino group of a lysine residue on the target protein (Hershko and Ciechanover 1998). In human cells, this latter reaction is catalyzed by one of the nearly 600 E3 ubiquitin ligases encoded by in the genome (Zheng and Shabek 2017). Similar E1-E2-E3 enzymatic cascades exist for the conjugation of the other Ubl modifiers although, compared to ubiquitin, their repertoire of E3 ligases is much more restricted, ranging from ∼8 E3s for SUMO to a single E3 for UFM1 (Cappadocia and Lima 2018; Vertegaal 2022).

There is considerable crosstalk and interaction among the various ubiquitin and Ubl conjugation systems. For example, there are a number of SUMO-targeted E3 ubiquitin ligases such as RNF4 that catalyze the conjugation of ubiquitin chains on SUMOylated substrates (Tatham et al. 2008; Chang et al. 2021). Another example of crosstalk germane to this work is the conjugation of the Ubl NEDD8 on the cullin subunit of the cullin-RING-ligase (CRL) complexes, which form a large fraction of the E3 ubiquitin ligase complement of eukaryotic cells (Harper and Schulman 2021). Site-specific NEDD8 conjugation (neddylation) on cullin proteins is necessary to activate the E3 ligase activity of CRLs and, conversely, inhibition of neddylation blocks CRL activity and leads to the stabilization of their substrates (Harper and Schulman 2021).

The control of genome stability is a process that is heavily influenced by degradative and non-degradative ubiquitylation, as well as SUMOylation (Jackson and Durocher 2013). For example, DNA double-strand breaks (DSBs) initiate a ubiquitin-dependent modification cascade on histones surrounding break sites, culminating with RNF168 ubiquitylating histone H2A at its N-terminus (Gatti et al. 2012; Mattiroli et al. 2012), a histone modification that is specifically read by the 53BP1 and BARD1 proteins that control DNA repair (Fradet-Turcotte et al. 2013; Wilson et al. 2016; Becker et al. 2021). In contrast, investigations on the role of SUMO in DNA repair by homologous recombination led to the concept that SUMO acts via group modification of target proteins, where multiple components of a pathway are SUMOylated to strengthen their collective action (Psakhye and Jentsch 2012; Vertegaal 2022). Additional Ubl proteins are also linked to genome maintenance processes, with ISG15 modification being identified by multiple groups as acting in the response to DNA replication stress (Park et al. 2014; Raso et al. 2020; Wardlaw and Petrini 2022).

The central role of ubiquitin-dependent degradation in controlling the activity of oncogenes and tumor suppressors have made the ubiquitin proteasome system (UPS) and Ubl conjugation pathways attractive drug targets. The regulatory approval of the proteasome inhibitor bortezomib for the treatment of multiple myeloma provided an important proof-of-concept of the clinical utility of targeting the UPS (Adams 2001). This spurred the development of various inhibitors that exploit druggable nodes in the UPS pathways. These include the E1 activating enzymes (Wertz and Wang 2019) as well as p97/VCP, a hexameric ATPase that “extracts” ubiquitylated substrates from membranes, protein assemblies and chromatin to facilitate their degradation (Meyer and Weihl 2014). There is also considerable interest in the development of agents that target the reversal of ubiquitin or Ubl conjugation such as inhibitors of deubiquitylating enzymes or of the COP9/signalosome that catalyzes de-neddylation (Wertz and Wang 2019).

In an effort to assess the function of ubiquitin and Ubl modifiers from the vantage point of cellular fitness, we sought to define how cells respond to perturbations in ubiquitin and Ubl conjugation pathways. We undertook genome-scale CRISPR Cas9 screens to identify genes that contribute to the cellular resistance to inhibition of ubiquitin, SUMO and NEDD8 conjugation as well as mapping genes that contribute to the response to p97/VCP inhibition. From these screens, we validated that the transmembrane proteins TMED2/TMED10 contribute to survival in response to p97/VCP inhibition, and also characterized the role of NFATC2IP, a protein with tandem SUMO-like domains, in promoting SUMO-dependent genome maintenance via a functional interaction with the SMC5/SMC6 complex. We contend that this work will provide a useful dataset to study the cellular functions of ubiquitin, SUMO and NEDD8, and identifies NFATC2IP as a key mediator of the role of SUMOylation in controlling genome integrity.

## Results

### A chemogenetic map of Ub/Ubl pathway inhibitors

In a recent chemogenetic survey of the response to genotoxic agents undertaken by our group (Olivieri et al. 2020), we screened for genes that modulate sensitivity to MLN4924, an inhibitor of the NEDD8 E1, since neddylation inhibition causes DNA damage (Soucy et al. 2009). To expand this dataset with the view of charting the genetic interactions underlying the cellular responses to inhibitors of ubiquitin (Ub) and Ubl conjugation pathways, we undertook additional chemogenomic CRISPR screens with the UBA1 (ubiquitin E1) inhibitor TAK-243 (Hyer et al. 2018), the SAE1 (SUMO E1) inhibitor TAK-981 (Langston et al. 2021), and the p97/VCP segregase inhibitor CB-5083 (Zhou et al. 2015). The screens were carried out at doses that killed ∼20% of cells (LD_20_) in the hTERT-immortalized retinal pigment epithelial-1 (RPE1-hTERT) *TP53^-/-^* cells stably expressing Cas9 (Zimmermann et al. 2018) as depicted in Figure S1A and as described previously (Olivieri et al. 2020; Olivieri and Durocher 2021). Gene-level normalized Z-scores (NormZ) were computed using DrugZ (Colic et al. 2019) and are shown in Figure 1A and Table S1.

**Figure 1.**
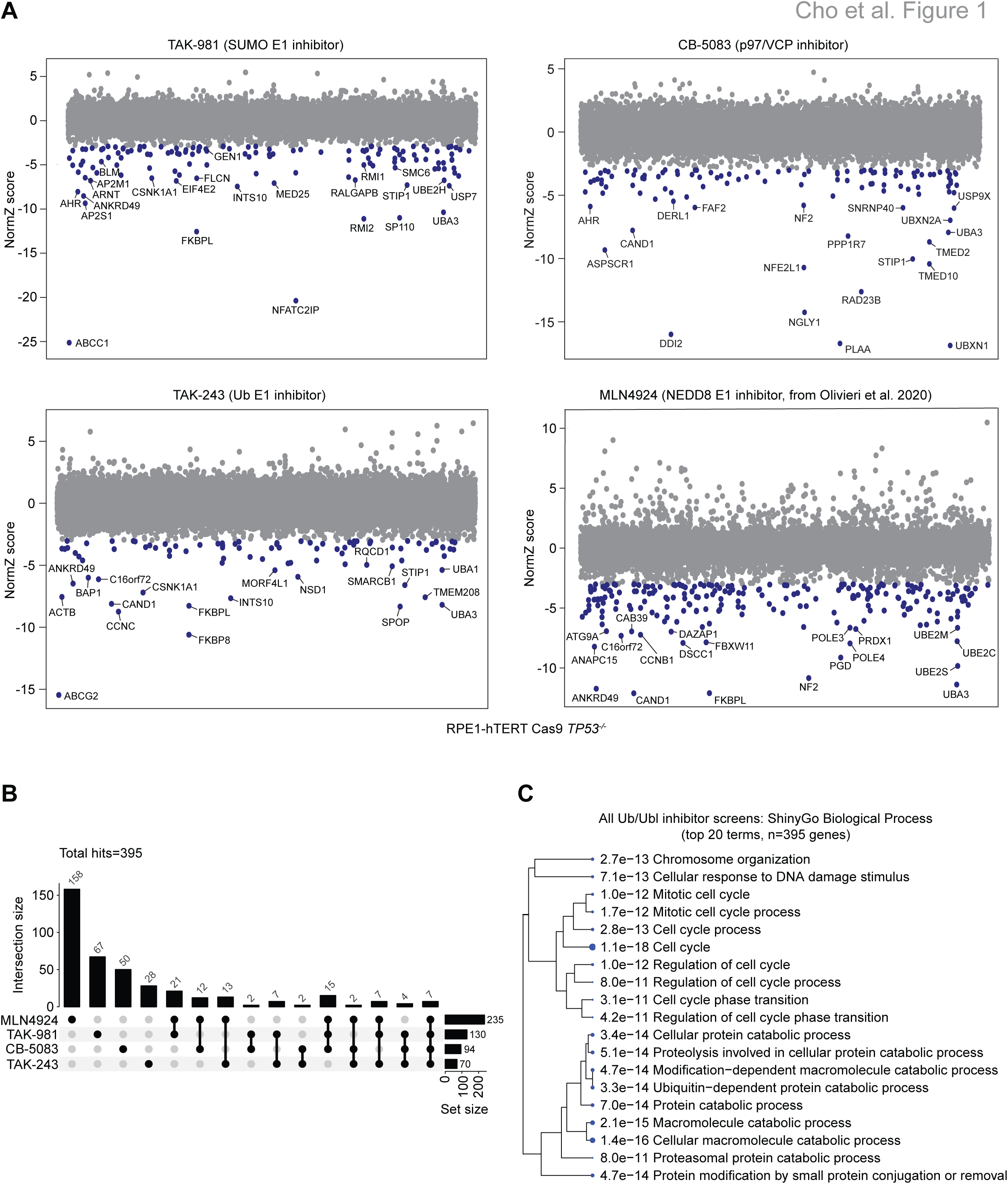
Chemogenetic CRISPR screens charting Ub/Ubl pathways. (A) Chemogenomic CRISPR screen results for RPE1-hTERT Cas9 *TP5*3^-/-^ cells treated with the indicated ubiquitin or Ubl conjugation pathway inhibitor. Each data point represents a gene score (normZ value) for an individual gene in the indicated inhibitor screen, calculated using DrugZ. Blue points represent sensitizing genes (normZ < -3 with FDR < 0.15). (B) UpSet plot summarizing the CRISPR screen results. The set size values represent the total number of sensitizing (normZ < -3) and resistance (normZ > 6) genes in each screen. (C) Gene ontology enrichment of biological process terms using ShinyGo for the 395 hits identified in the four screens. An FDR cut-off of 0.05 was used, and the top 20 enriched functional terms are shown with their individual *P*-values. The relative size of the circle for each GO term represents significance of *P-*value.

To remain consistent with the analyses of Olivieri et al. (2020), we selected NormZ values less than -3 with false discovery rates (FDR) lower than 15% to identify genes whose mutation caused sensitization to the inhibitors. For genes whose mutation caused resistance to inhibitor treatment, we selected a NormZ value greater than 6. These cut-offs identified a total of 395 genes that modulate the response of RPE1-hTERT cells to drug-mediated inhibition of Ub/Ubl pathways, with 92 genes being identified in two screens or more. We identified 235 hits for the MLN4924 screen, 130 for TAK-981, 94 for CB-5083, and 70 for TAK-243, (Figure 1B and Tables S1). We analyzed the connectivity of the 395 genes by building networks based on protein-protein interactions, using STRING (Figure S1B and Table S2) (Szklarczyk et al. 2021) or gene-gene essentiality score correlations in DepMap (Figure S2 and Table S2) (Dempster et al. 2019). When the 395 genes were mapped onto either the BioGRID or CORUM protein interaction datasets, 372 or 136 proteins encoded by genes in our hit gene set, respectively, were physically connected to at least one other protein in the dataset, which represents a statistically significant enrichment in protein-protein interactions (Oughtred et al. 2021; Tsitsiridis et al. 2023) (Figure S3A). A number of distinct submodules were apparent in the protein-protein interaction network, including a module surrounding p97/VCP that was connected to a second submodule comprising HSP90 and HSP70 chaperones and adaptor proteins (Figure S1B). At the Pearson correlation coefficient threshold used (0.25), the DepMap-based network presented a highly connected network of 240 genes with some clear functional subnetworks (Figure S2). As an example, multiple genes encoding factors involved in protein glycosylation such as *B4GALT7*, *SLC35B2* and *NDST1* formed a clear subcluster of genes whose disruption was primarily causing sensitivity to MLN4924, suggesting that defects in protein glycosylation imposes a required response for CRL E3 ligases or non-cullin neddylation, possibly through the unfolded protein response.

Functional term analysis using ShinyGO (Ge et al. 2020) for the 395 genes using gene ontology (GO) Biological Process (BP) showed enrichment in pathways regulated by ubiquitin or Ubl modification, and in pathways pertaining to modifications by ubiquitin or Ubl proteins (Figure 1C; GO term enrichment for individual screens are in Figure S3B). For example, “ubiquitin-dependent protein catabolic process” (GO:0006511) was highly enriched (p=3.3 × 10^-14^) alongside processes known to be modulated by ubiquitin or Ubls such as regulation of “mitotic cell cycle” (GO:0000278; 1.0 × 10^-12^) or “cellular response to DNA damage stimulus” (GO:0006974; 7.1 × 10^-13^). From these analyses, we conclude that the screens were successful in probing pathways that are relevant to ubiquitin and Ubl biology.

### *TMED2-TMED10* promotes resistance to p97 inhibition

To assess the usefulness of the dataset in providing new biological insights, we first examined the *TMED2* and *TMED10* genes that were among the top hits in the CB-5083 screen (Figure 1A and Table S1), which also identified genes encoding multiple p97/VCP adaptor proteins, UBXN1, UBXN2A, ASPSCR1, and PLAA (Ye et al. 2017), as well as proteins involved in NRF1 (encoded by *NFE2L1*) activation and processing such as *NGLY1*, *DDI2* and *NFE2L1* itself (Figure 1A). Both TMED2 and TMED10 are single-pass transmembrane proteins found in the Golgi and endoplasmic reticulum (ER) where they participate in the transport of GPI-anchored protein (Bonnon et al. 2010). We were attracted by these proteins since NRF1 is an ER-regulated transcription factor that controls proteasome subunit gene transcription, and whose activation is dependent on p97/VCP, which acts as part of a complex proteolytic processing event that causes NRF1 to translocate from the ER to the cytoplasm and then to the nucleus (Radhakrishnan et al. 2014; Northrop et al. 2020; Ruvkun and Lehrbach 2023). Given the presence of TMED2/TMED10 in the Golgi and ER, along with their known role in protein transport, we initially hypothesized that these proteins may participate in NRF1 activation.

We first validated that two independent sgRNAs targeting *TMED2* and *TMED10* sensitize cells to CB-5083 treatment by using clonogenic survival assays that employed sgRNAs targeting *DDI2* as a positive control (Figure 2A,B). We next examined if the loss of TMED2/10 influenced NRF1 processing by assessing NRF1 levels and isoforms by immunoblotting. Under normal conditions, NRF1 is synthesized as an ER-localized transmembrane protein but is then retrotranslocated into the cytoplasm where it is rapidly degraded by the proteasome (Steffen et al. 2010; Radhakrishnan et al. 2014). However, under conditions of limiting proteasome activity (such as in response to proteasome inhibition), NRF1 is proteolytically processed by DDI2 to produce an isoform competent for nuclear translocation and transcriptional activation (Northrop et al. 2020). To our surprise, loss of either TMED2 or TMED10 led to increased, rather than decreased, levels of all NRF1 isoforms in response to proteasome inhibition with carfilzomib (Demo et al. 2007) (Figure 2C). This increase in NRF1 was accompanied by an increase in *PSMB7* and *PSMC4* mRNA levels, suggesting higher NRF1 activity (Figure 2D). Contrary to our initial expectation, these observations indicate that TMED2/10 are unlikely to promote resistance to p97/VCP inhibition by promoting NRF1 processing, and thus we suspect that TMED2/10 could impact ER-associated quality control pathways in a manner that imposes a higher burden on p97/VCP activity. In support of this possibility, we note that additional genes encoding proteins involved in such processes (*SEC63* and *DERL1*) were also identified as genes promoting resistance to CB-5083 (Figure 1A and Table S1).

**Figure 2.**
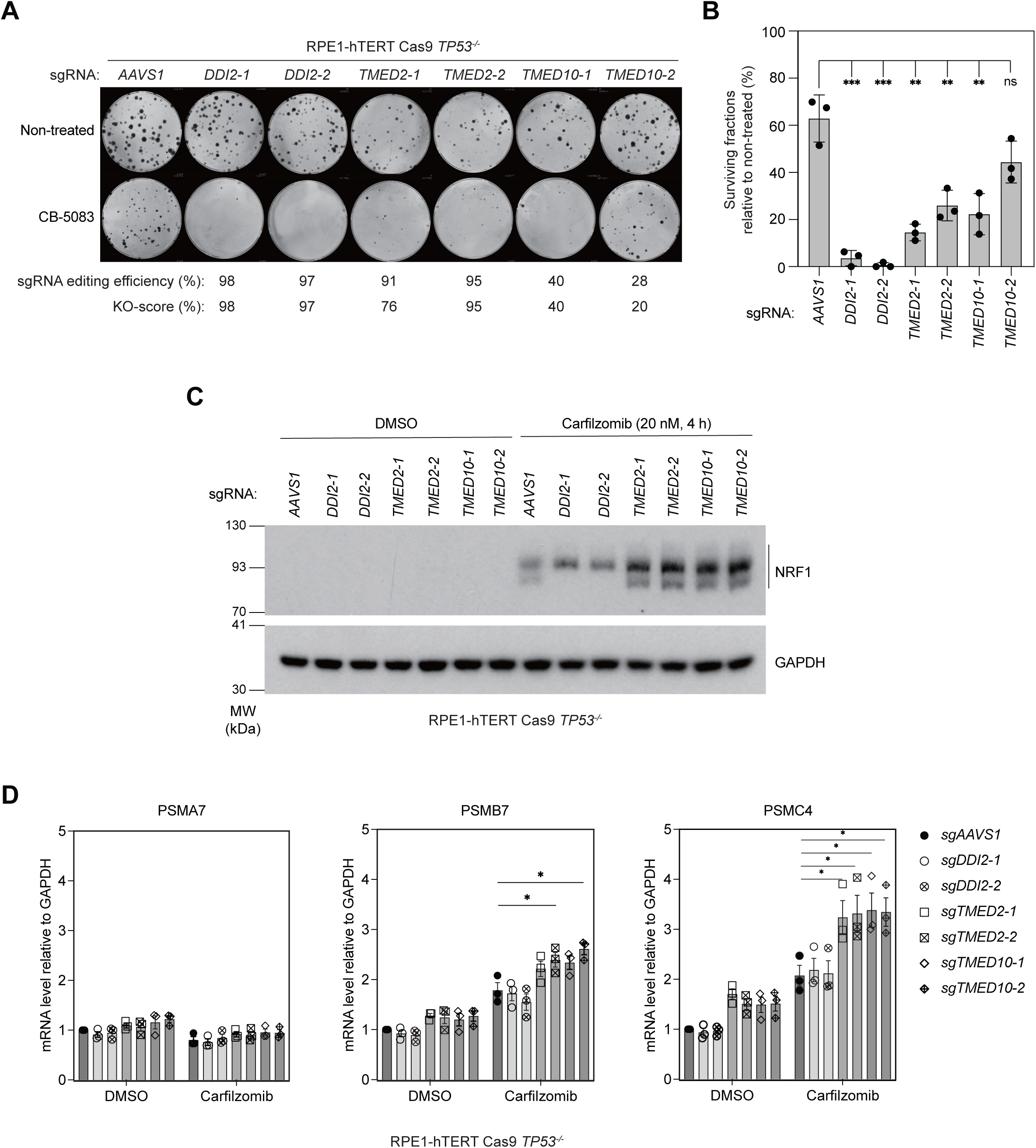
*TMED2*/*TMED10* promotes resistance to p97 inhibitor CB-5083. (A) and (B) Clonogenic survival assays of RPE1-hTERT Cas9 *TP53^-/-^* cells transduced with a virus expressing the indicated sgRNA in response to p97 inhibitor CB-5083 treatment (200 nM). (A) Representative images of clonogenic survival assay plates. Percentage of the total cell population that underwent CRISPR-mediated gene editing (sgRNA editing efficiency) and percentage of the cell population that yielded an out-of-frame gene edit (KO-score) from each sgRNA are indicated. (B) Quantitation of the experiment shown in (A). Percentage of surviving fractions were normalized to the non-treated (DMSO) condition for each sgRNA tested. The bars represent the mean ± s.e.m. (n=3). Statistical comparisons were made to the surviving fraction after transduction with sg*AAVS1* by performing an unpaired t-test. ***: *P* < 0.001. **: *P* < 0.01. ns: *P ≥* 0.05. (C) Immunoblot analysis of NRF1 expression in RPE1-hTERT Cas9 *TP53^-/-^* cells transduced with the indicated sgRNA and treated with carfilzomib (20 nM) for 4 h or left untreated. GAPDH was used as a loading control. (D) Quantitative RT-PCR to detect mRNA of PSMA7, PSMB7, and PSMC4 using TaqMan assays (Table S4) from extracts of RPE1-hTERT Cas9 *TP53^-/-^* cells treated with carfilzomib as in (C). Bars represent the mean ± s.e.m. (n=3). Statistical comparisons were made to control condition from the sg*AAVS1*-transduced cells by performing an unpaired t-test. *: *P* < 0.05.

### *NFATC2IP* promotes survival in response to SUMOylation inhibition

Another gene that attracted our attention was *NFATC2IP* (also known as *NIP45*) as this gene ranked just behind that encoding the multidrug resistance transporter MRP1 (*ABCC1*) (Robey et al. 2018), in the TAK-981 screen (Figure 1A and Table S1). *NFATC2IP* encodes a protein featuring two SUMO-like domains (SLDs; Figure 3A) and is the likely ortholog of the *Saccharomyces cerevisiae* Esc2 and *Schizosaccaromyces pombe* Rad60 proteins (Novatchkova et al. 2005) (Figure 3A). In yeast species, Esc2 and Rad60 promote replication fork integrity and tolerance to replication stress (Morishita et al. 2002; Boddy et al. 2003; Miyabe et al. 2006), but surprisingly in human cells, NFATC2IP has only been described as a co-factor of the nuclear factor of activated T-cells, cytoplasmic 2 (NFATc2) transcription factor (Hodge et al. 1996). Since NFATc2 was not a hit in the TAK-981 screen (Table S1) and since the TAK-981 screen hits were instead enriched for genes acting in the response to DNA damage (such as *BLM* and *RMI1/2*; Figure 1A and Figure S3B), we asked whether NFATC2IP promoted the normal cellular resistance to SUMOylation inhibition through a role in genome maintenance.

**Figure 3.**
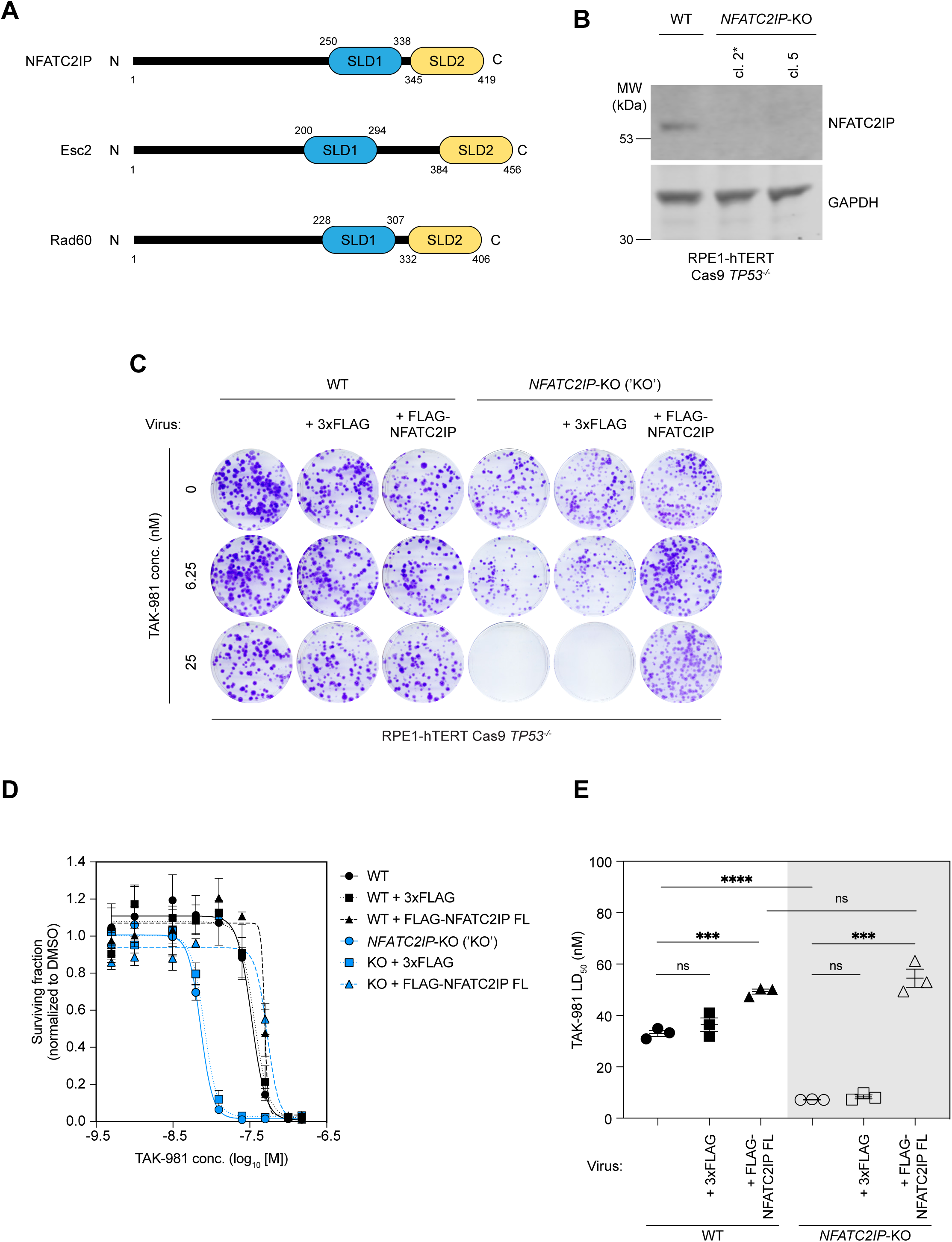
*NFATC2IP* promotes survival in response to SUMOylation inhibition. (A) Schematic of the structural domains of the human NFATC2IP protein and its ortholog proteins of the yeasts *Saccharomyces cerevisiae* (Esc2) and *Schizosaccharomyces pombe* (Rad60). SLD1: SUMO-like domain 1, SLD2: SUMO-like domain 2. (B) Immunoblot analyses of clonal *NFATC2IP-*KO cell lines isolated from a pool of RPE1-hTERT Cas9 *TP53^-/-^* cells transfected with a Cas9 ribonucleoprotein complex containing *NFATC2IP*-targeting sgRNA2 (Table S4). GAPDH was used as a loading control. Asterisk (*) indicates the *NFATC2IP*-KO clone that was used for all further experiments in this study. (C), (D), and (E) TAK-981 dose-response clonogenic survival assays in parental RPE1-hTERT Cas9 *TP53^-/-^* (WT) or isogenic *NFATC2IP*-KO (KO) cells that were either transduced with lentivirus expressing 3xFLAG-tagged NFATC2IP, 3xFLAG alone, or left untransduced. (C) Representative images of the clonogenic survival assay. (D) Quantitation of surviving fractions normalized to the DMSO vehicle condition for the experiment shown in (C). Data are shown as the mean ± s.e.m. (n=3). FL: Full-length. (E) Determination of TAK-981 LD_50_. Data is shown as the mean ± s.e.m. (n=3). Statistical comparisons were performed using two-tailed unpaired t-tests. ****: *P* < 0.0001. ***: *P* < 0.001. ns: *P ≥* 0.05. FL: Full-length.

We first generated clonal knock-outs (KO) of *NFATC2IP* in RPE1-hTERT *TP53^-/-^* Cas9 cells (Figure 3B and Figure S4A,B) with CRISPR gene editing that generated a biallelic one-nucleotide deletion (c.493delC), which caused a frameshift mutation (p.His165MetfsX15). Using these clonal *NFATC2IP*-KO cells, we assessed the half-maximal lethal dose (LD_50_) of TAK-981 in clonogenic survival assays. Loss of NFATC2IP decreased the TAK-981 LD_50_ from 28.5 ± 2.5 nM, in the parental cell line, to 7.0 ± 0.8 nM (Figure 3C-E). This hypersensitivity of *NFATC2IP*-KO cells was fully reversed by re-introducing exogenous NFATC2IP expressed as 3xFlag-tagged protein from a lentiviral vector (WT; Figure 3C-E and Figure S4C). We conclude that NFATC2IP promotes cell survival upon SUMOylation inhibition.

### NFATC2IP promotes genomic integrity

Given the role of NFATC2IP orthologs in genome maintenance, as well as the central role of SUMO in protecting genome integrity, we tested whether loss of NFATC2IP caused genome instability. We monitored micronucleation, a sensitive readout of genome instability (Fenech et al. 2011), in parental and *NFATC2IP*-KO cells, with or without TAK-981 treatment using an automated microscopy pipeline. When SUMOylation is unperturbed, loss of NFATC2IP did not impact micronucleation levels (Figure 4A,B). However, following SUMO E1 inhibition, *NFATC2IP*-KO cells displayed enhanced micronuclei (MN) formation at all doses, and this phenotype was entirely suppressed by re-expression of NFATC2IP tagged either with 3xFlag or GFP (Figure 4A,B and Figure S5A-D). NFATC2IP therefore suppresses the genome instability caused by inhibition of SUMOylation.

**Figure 4.**
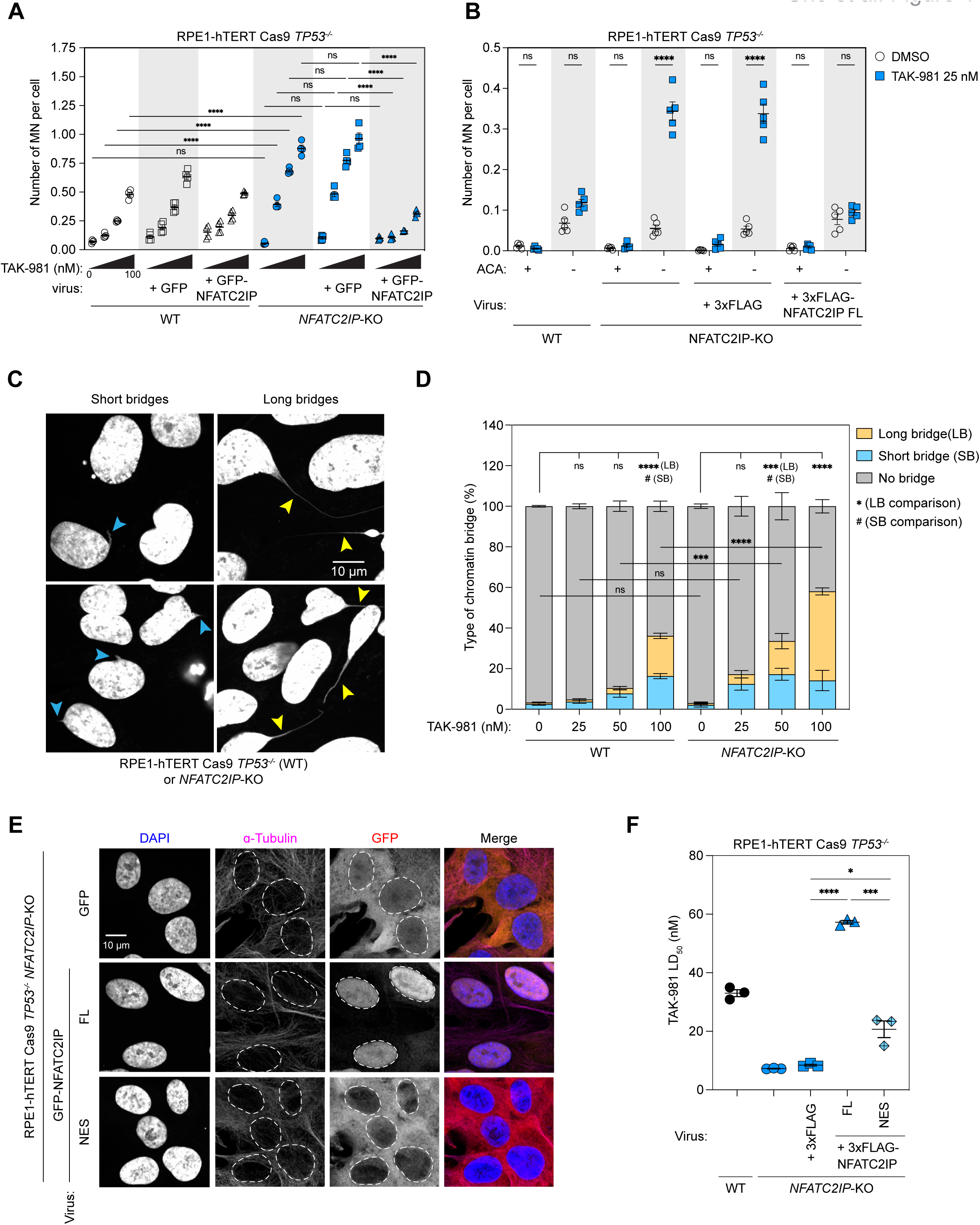
NFATC2IP protects genome integrity when SUMOylation is disrupted. (A) Quantitation of MN formation after treatment with increasing doses of TAK-981 for 72 h in RPE-hTERT Cas9 *TP53^-/-^* (WT) or isogenic *NFATC2IP*-KO cells that were either transduced with the indicated lentivirus or left untransduced. Data are shown as the number of MN per cell. A minimum of 455 nuclei were counted for each condition. Bars represent the mean ± s.e.m. (n=4). Multiple unpaired t-tests were used for statistical analyses with Bonferroni-Dunn correction. ****: *P* < 0.0001. ns: *P ≥* 0.05. (B) Quantitation of ACA-positive or -negative MN formation after treatment with or without TAK-981 (25 nM for 48 h) in parental RPE1-hTERT Cas9 *TP53^-/-^* (WT) or isogenic *NFATC2IP*-KO cells that were either transduced with the indicated lentivirus or left untransduced. Data are shown as the number of MN per cell, and a minimum of 273 nuclei were counted for each condition. Bars represent the mean ± s.e.m. (n=5). FL: Full-length. Multiple unpaired t-tests were used for analyses with Bonferroni-Dunn correction. ****: *P* < 0.0001. ns: *P ≥* 0.05. (C) Representative micrographs of chromatin bridge formations visualized by DAPI staining. Blue arrowheads point to chromatin bridges that were identified as short bridges (< ∼15 µm), and yellow arrowheads point to long chromatin bridges (> ∼15 µm). Scale bar=10 µm. (D) Quantitation of nuclei displaying chromatin bridges following treatment with the indicated doses of TAK-981 for 24 h in RPE1-hTERT Cas9 *TP53^-/-^* (WT) or isogenic *NFATC2IP*-KO cells. Data are shown as the percentage of nuclei counted that contain either short or long chromatin bridges. A minimum of 263 nuclei were assessed for each condition. Bars represent the mean ± s.e.m. (n=3). Analyses were performed using two-way ANOVA with Tukey’s multiple comparison testing. For comparisons of long chromatin bridges, ****: *P* < 0.0001. ***: *P* < 0.001. ns: *P ≥* 0.05. For comparisons of short chromatin bridges, only significant comparisons were shown. #: *P* < 0.05. LB: long bridge. SB: short bridge. (E) Representative immunofluorescence micrographs assessing the localization of GFP-tagged NFATC2IP in RPE1-hTERT Cas9 *TP53^-/-^ NFATC2IP*-KO cells. Cells were stained with DAPI as a nuclear marker and antibodies against the indicated proteins. FL: Full-length. NES: NES-NFATC2IP. Scale bar=10 µm. (F) TAK-981 LD50 values of independent clonogenic survival assays in parental RPE1-hTERT Cas9 *TP53^-/-^*(WT) or isogenic *NFATC2IP*-KO, or *NFATC2IP*-KO cells that were left untransduced or were transduced with a lentivirus encoding the indicated protein. Data is shown as the the mean ± s.e.m. (n=3). FL: full-length. NES: NES-NFATC2IP. Statistical comparisons were performed using two-tailed unpaired t-tests. ****: *P* < 0.0001. ***: *P* < 0.001. *: *P* < 0.05.

MN formation can occur via two broadly distinct routes: either through whole chromosome mis-segregation, or through the segregation of acentric fragments (Fenech et al. 2011). It is possible to distinguish between these possibilities simply by monitoring kinetochores and centromeres using anti-centromere antibodies (ACA). We observed that upon TAK-981 treatment, *NFATC2IP*-KO cells produce primarily centromere-negative (ACA-) MN (Figure 4B). These results suggest that NFATC2IP guards against acentric chromosome mis-segregation when SUMOylation is impaired.

In addition to micronucleation, we observed that the combined loss of NFATC2IP and TAK-981 treatment was accompanied by the formation of long chromatin bridges connecting two nuclei (Figure 4C). We quantitated the formation of DAPI-stained chromatin bridges and binned the data according to whether we observed cells with normal chromatin or that displayed either short or long chromatin bridges (i.e. bridges > 15 μm). We observed that loss of NFATC2IP led to an increase in long chromatin bridges following TAK-981 treatment at all doses tested compared to the parental cell line, with the strongest effect seen at the 50 nM TAK-981 dose (Figure 4D). As with micronucleation, SUMOylation inhibition on its own is able to induce chromatin bridge formation but *NFATC2IP*-KO cells accumulate bridges at higher frequency and at lower concentrations of TAK-981 than wild type cells, indicating that the loss of NFATC2IP exacerbates a pathological process induced by inhibition of the SUMO E1 (Figure 4D). Since breakage of chromatin bridges during cytokinesis can generate acentric fragments (Warecki and Sullivan 2020; Hong et al. 2021), we surmise that they may be a source for the observed micronucleation and, possibly, the hypersensitivity of *NFATC2IP*-KO cells to SUMO E1 inhibition.

### NFATC2IP acts in interphase to promote genome maintenance

Chromatin bridges can originate either from defective DNA replication, repair, recombination or via the failure to complete mitotic processes like chromosome decatenation (Hong et al. 2021). To begin distinguishing between these possibilities, we asked whether NFATC2IP acted to promote genome integrity in interphase or during mitosis. To do so, we fused a nuclear export signal (NES) to NFATC2IP so that it can only access chromatin after nuclear envelope breakdown at the onset of mitosis. As expected, the NES-NFATC2IP proteins, expressed either as GFP or 3xFlag fusions, were restricted to the cytoplasm during interphase (Figure 4E and Figure S5E). The nucleus-excluded form of NFATC2IP failed to fully restore resistance to TAK-981 in *NFATC2IP*-KO cells, unlike its wild type counterpart (FL; Figure 4F). These observations indicate that NFATC2IP acts in interphase, requiring access to chromatin prior to mitosis to promote genome integrity when SUMOylation is impaired.

### Structure-function analysis of NFATC2IP

To gain insights into the mechanism by which NFATC2IP may promote genomic integrity when SUMOylation is perturbed, we assessed the involvement of the SLD2 domain (Figure 3A) in promoting cell survival in response to TAK-981 treatment given the role of this domain in Esc2 and Rad60 (Prudden et al. 2009; Sebesta et al. 2017; Li et al. 2021). We expressed, in *NFATC2IP*-KO cells, NFATC2IP lacking the SLD2 domain (ΔSLD2) or a variant that harbored the D394R mutation that corresponds to Rad60 E380R, a point mutation that abolishes interaction with the SUMO E2 UBC9 (Prudden et al. 2009; Sekiyama et al. 2010; Prudden et al. 2011). We found that loss of SLD2 or disruption of the putative interaction with UBC9 failed to restore normal resistance to TAK-981 (Figure 5A and Figure S6A,B), suggesting that the NFATC2IP SLD2 domain is critical for promoting survival in response to SUMOylation inhibition. Similarly, while micronucleation after TAK-981 treatment in NFATC2IP-deficient cells could be completely rescued by expression of epitope-tagged NFATC2IP, neither NFATC2IP D394R nor ΔSLD2 could do so (Figure 5B and Figure S6B). We also tested whether expression of NFATC2IP ΔSLD1 in *NFATC2IP*-KO cells could suppress TAK-981-induced micronucleation and found it did so partially (Figure 5B and Figure S6B). Together, these results indicate that NFATC2IP promotes genome integrity and cellular survival in response to SUMO E1 inhibition in a manner that requires its SLD2 domain, with some contribution from SLD1.

**Figure 5.**
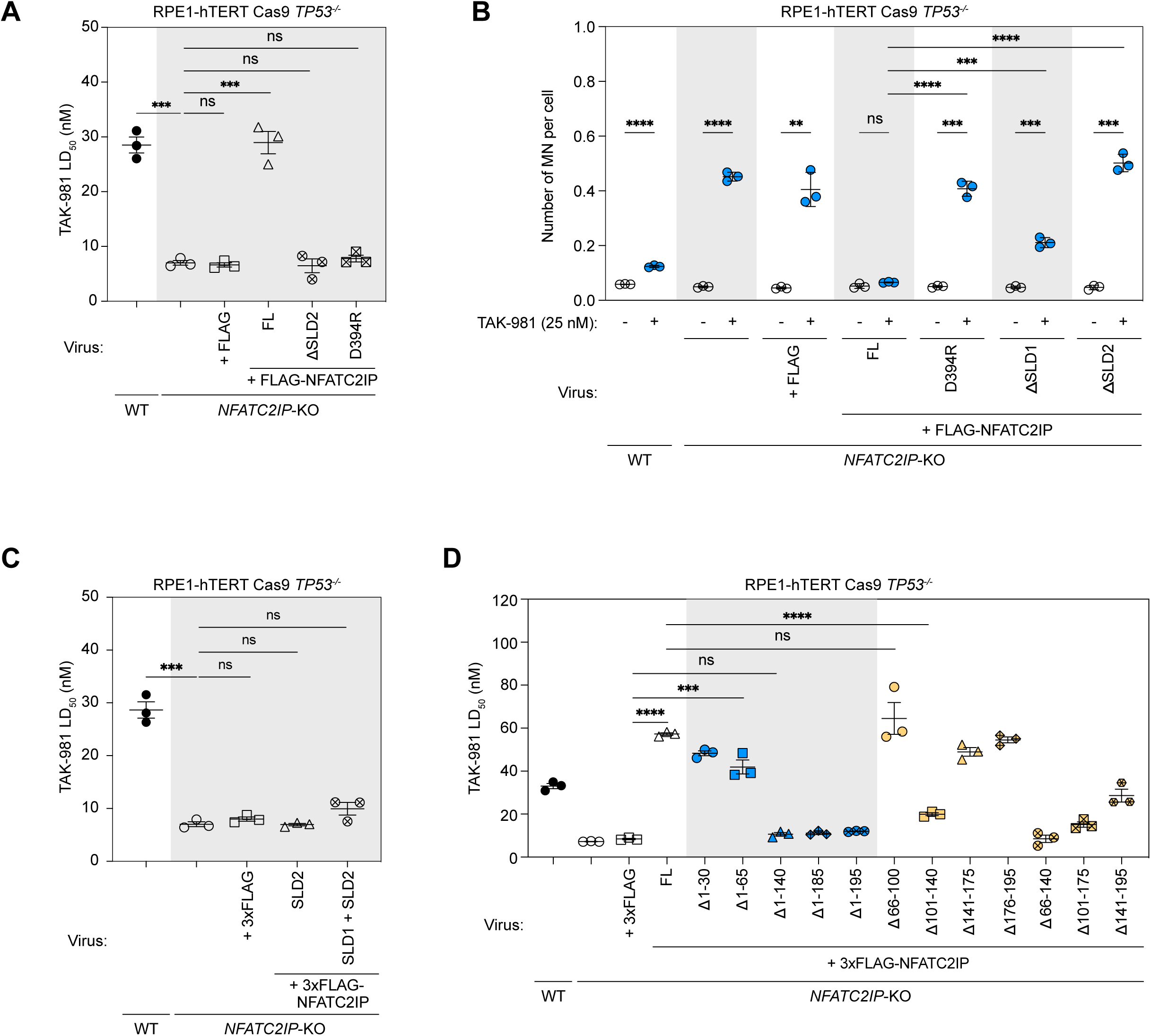
Structure-function analysis of NFATC2IP. (A) –(D) RPE1-hTERT Cas9 *TP53^-/-^* (WT) or isogenic *NFATC2IP*-KO cells were either left untransduced or were transduced with lentivirus encoding the indicated protein. FL: full-length. (A) Determination of TAK-981 LD_50_. Data are shown as the mean ± s.e.m. (n=3). Statistical comparisons were with two-tailed unpaired t-tests. ***: *P* < 0.001. ns: *P ≥* 0.05. ΔSLD2: deletion of amino acid residue 345–419. (B) Quantitation of MN formation after treatment with TAK-981 (25 nM) for 48 h. Data are shown as the number of MN per cell, and a minimum of 1229 nuclei were counted for each condition. Bars represent the mean ± s.d. (n=3). Data were analyzed with multiple unpaired t-test with Bonferroni-Dunn correction. ****: *P* < 0.0001. ***: *P* < 0.001. **: *P* < 0.01. ns: *P ≥* 0.05. ΔSLD1: deletion of amino acid residue 256–344. ΔSLD2: deletion of amino acid residue 345– 419. (C) Determination of TAK-981 LD_50_. Data are shown as the mean ± s.e.m. (n=3) and were analyzed with two-tailed unpaired t-tests. ***: *P* < 0.001. ns: *P ≥* 0.05. SLD2: NFATC2IP amino acid residues 345–419. SLD1 + 2: NFATC2IP amino acid residues 251–419. (D) Determination of TAK-981 LD_50_. Data are shown as the mean ± s.e.m. (n=3) and were analyzed with two-tailed unpaired t-tests. ****: *P* < 0.0001. ***: *P* < 0.001. ns: *P ≥* 0.05. Deletion mutants of NFATC2IP are schematized in Figure S7.

Despite the functional importance of the SUMO-like domains, we found that expression of SLD2 or a fragment encompassing SLD1-SLD2 were insufficient to restore resistance of *NFATC2IP*-KO cells to SUMO E1 inhibition (Figure 5C and Figure S6CD) indicating that additional segments of the NFATC2IP protein contribute to its function. Therefore, to further dissect the structure-function relationship of NFATC2IP, we expressed protein deletion mutants (schematized in Figure S7) in *NFATC2IP*-KO cells and calculated their TAK-981 LD_50_ in clonogenic survival assays. From this analysis, we identified an additional region that contributes to NFATC2IP function in response to inhibition of SUMOylation that is encompassed within residues 101–140 (Figure 5D and Figure S6E-H). Furthermore, our data suggests that most of the first 100 amino acid residues of NFATC2IP are dispensable for its role in promoting resistance to TAK-981.

### NFATC2IP interacts with the SMC5/SMC6 complex

We were struck by the similarities between the chromosome segregation phenotypes of *NFATC2IP*-KO cells following TAK-981 treatment and those of *SMC5-*, *SMC6-* and *NSMCE2*-deficient cells (Gallego-Paez et al. 2014; Payne et al. 2014; Jacome et al. 2015; Pryzhkova and Jordan 2016). These genes encode for factors that constitute the SMC5/6 complex which ensures proper segregation of chromosomes and genome integrity, with NSMCE2 acting as a SUMO E3 ligase (Potts and Yu 2005; Aragón 2018; Venegas et al. 2020). Since Esc2/Rad60 is functionally linked with the SMC5/6 complex (Morishita et al. 2002; Sollier et al. 2009; Choi et al. 2010; Heideker et al. 2011; Prudden et al. 2011), we explored the possibility that NFATC2IP may mediate resistance to SUMOylation inhibition by collaborating with the SMC5/6 complex. To do so, we first surveyed the potential for protein-protein interactions between NFATC2IP and SMC5/6 complex members using AlphaFold-Multimer (AF-Multimer; Evans et al. 2021; Jumper et al. 2021; Mirdita et al. 2022) by testing pairwise combinations among SMC5/6 complex members, known SMC5/6 regulators that included SLF1, SLF2 and RAD18, and NFATC2IP as previously described (Sifri et al. 2023) (Table S3). This analysis recapitulated known protein-protein interactions in this complex, including the interaction between the BRCT domains of SLF1 and phosphorylatable residues in RAD18 (Raschle et al. 2015), or the NSMCE3 subunit with NSMCE1 and NSMCE4 (Figure 6A and Table S3) (Adamus et al. 2020; Vondrova et al. 2020; Yu et al. 2021; Yu et al. 2022).With respect to NFATC2IP, AF-Multimer predicted an interaction with the SMC5 subunit of the SMC5/6 complex and the NFATC2IP SLD1 domain (Figure 6A,B and Table S3). We validated these predictions in co-immunoprecipitation studies in 293T cells with Flag-tagged NFATC2IP variants and endogenous SMC5, which showed that the NFATC2IP-SMC5 interaction was absolutely dependent on SLD1, and to a lesser extent, SLD2 (Figure 6C). Furthermore, NFATC2IP interacts with UBC9 via its SLD2 (Prudden et al. 2009; Prudden et al. 2011), raising the intriguing possibility that NFATC2IP makes contact with the SMC5/6-associated SUMO E3 ligase and suggesting a model where NFATC2IP could position UBC9 near the NSMCE2 RING domain. We therefore carried out a new round of AF-Multimer predictions with NFATC2IP, UBC9, the coiled-coil region of SMC5 and NSMCE2 (Figure 6D). A robust structural model was computed that was consistent with NFATC2IP SLD2 interacting with NSMCE2 through simultaneous interactions with UBC9 via distinct interfaces (Figure 6D). Interestingly, the SLD2-UBC9 interaction occurs via the same “SUMO backside” site on UBC9 that promotes SUMOylation by yeast Nse2 (Figure 6E) (Varejao et al. 2021). Therefore, our interaction and modelling data suggest that NFATC2IP may act as a positive regulator of NSMCE2-dependent SUMOylation (Figure 6F).

**Figure 6.**
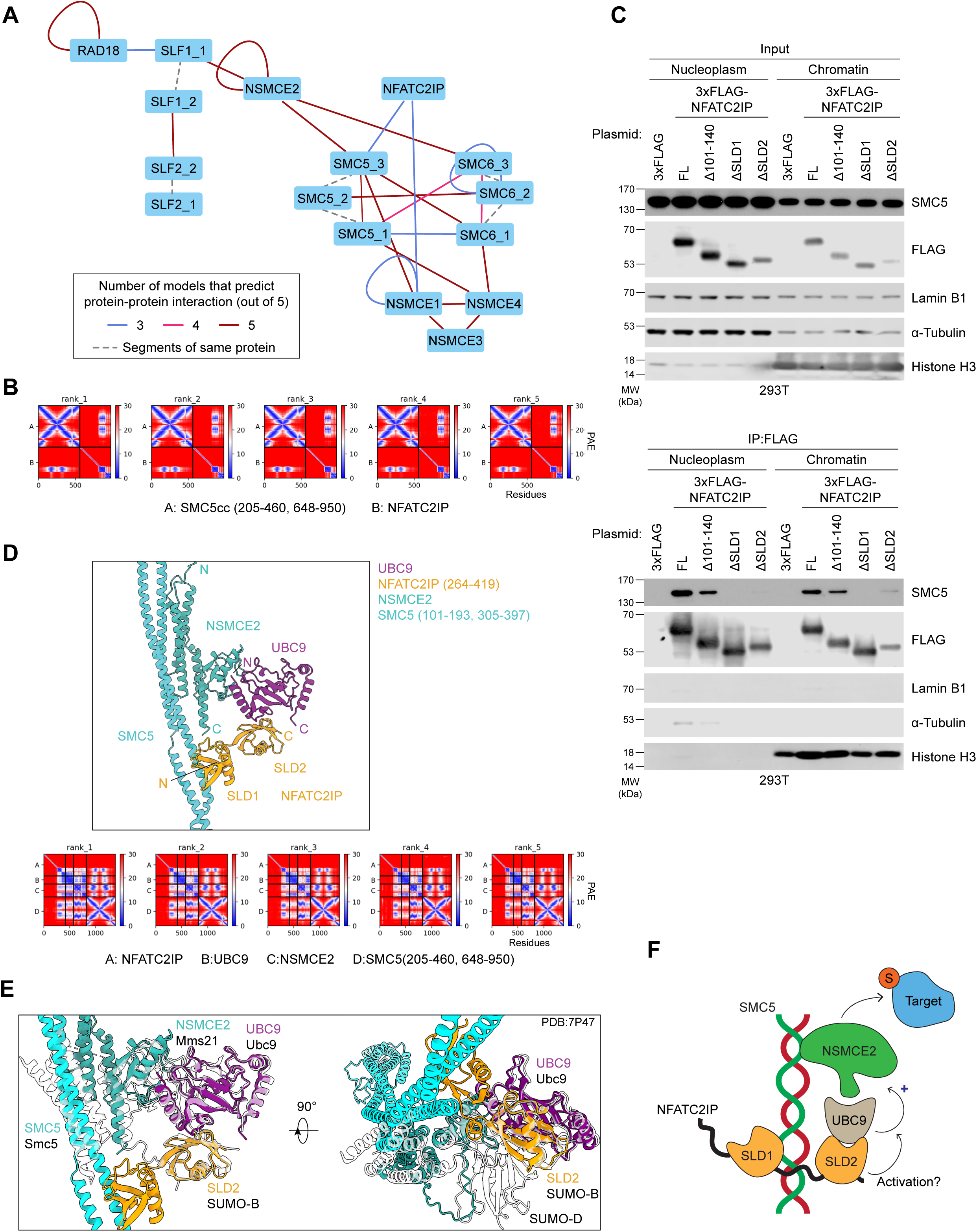
NFATC2IP binds the SMC5/SMC6 complex through its SUMO-like domains. (A) Schematic of pairwise matrix screens by AF-Multimer to predict protein-protein interactions among NFATC2IP and the SMC5/SMC6 complex subunits. Predicted interactions where at least 3 out of 5 models that met the cut-off scores of pDockQ < 0.23 and interface PAE < 15 Å are shown. Large proteins (SMC5, SMC6, SLF1, and SLF2) were divided into fragments according to either experimentally-determined or AF2-predicted structural domain boundaries. Fragments of the same protein are indicated with dashed lines connecting the nodes. For SMC5, SMC5_1: ‘head’ region, residues 1–204, 951–1101. SMC5_2: ‘hinge’ region, resides 461-647. SMC5_3: ‘coiled-coil’ region, resides 205–460, 648–950. For SMC6, SMC6_1: ‘head’ region, resides 1– 201, 952–1091. SMC6_2: ‘hinge’ region, residues 476–662. SMC6_3: ‘coiled-coil’ region, residues 202–475, 663–951 (derived from the human SMC5/6 complex structure (Adamus et al. 2020). For SLF1, SLF1_1: residues 1–364. SLF1_2: residues 365–1058. For SLF2, SLF2_1: residues 1–600. SLF2_2: residues 601–1173. (B) PAE plots of the interaction between the coiled-coil region of SMC5 (SMC5_3 fragment of the pairwise modeling) and NFATC2IP, ranked by predicted template model (pTM) scores. (C) Subcellular fractions of 293T cells transiently expressing the indicated Flag-tagged NFATC2IP constructs were subjected to immunoprecipitation with anti-Flag antibodies. Input samples and immunoprecipitation products were immunoblotted with the indicated antibodies. α-tubulin, lamin B1, or histone H3 were included as controls for cytoplasmic, nucleoplasmic, or chromatin subcellular fractionations, respectively. FL: full-length. (D) Top panel: representative model of the AF-Multimer predictions of protein-protein interactions among the NFATC2IP SLDs (yellow), NSMCE2 (sea green), UBC9 (purple), and a portion of the SMC5cc region (cyan). Bottom panel: the PAE plots of the prediction models associated with the top panel. (E) Overview of the representative AF-Multimer prediction model (colours as in (D)) overlaid with the structure of the Smc5/Nse2 complex with Ubc9∼SUMO mimetic (PDB: 7P47, translucent). Proteins in the crystal structure are indicated alongside the AF-Multimer model at corresponding positions. (F) Functional model of the NFATC2IP-SMC5/6-NSMCE2-UBC9 complex. Arrow indicates the potential positive regulation of NSMCE2-dependent SUMOylation by NFATC2IP.

### NFATC2IP promotes chromatin SUMOylation

To investigate whether loss of NFATC2IP affects the levels of SUMOylation in cells, parental and *NFATC2IP*-KO cells were transduced with a plasmid stably expressing either His_6_-SUMO1 or -SUMO2. SUMOylated proteins were purified from cell lysates using nickel-nitrilotriacetic acid (Ni-NTA) agarose beads in strong denaturing conditions, and global SUMOylation levels were monitored by immunoblotting with antibodies to SUMO1 (Figure 7A) or SUMO2/3 (Figure 7B). As control for the affinity purification, we monitored RanGAP1, which is a well-characterized SUMOylated protein (Matunis et al. 1996). We found that in contrast to the abundance of SUMO1-modified proteins, which showed no difference in the pulldowns from the lysates derived of parental (WT) and *NFATC2IP*-KO cells (Figure 7A), there was a noticeable decrease in the amount of high molecular-weight SUMO2/3-modified proteins pulled down from *NFATC2IP*-KO cells, compared to those purified from parental cells (Figure 7B). The same was seen for SUMO2/3-modified RanGAP1 (Figure 7B). These data suggested that NFATC2IP may promote SUMOylation of proteins primarily by SUMO2/3.

**Figure 7.**
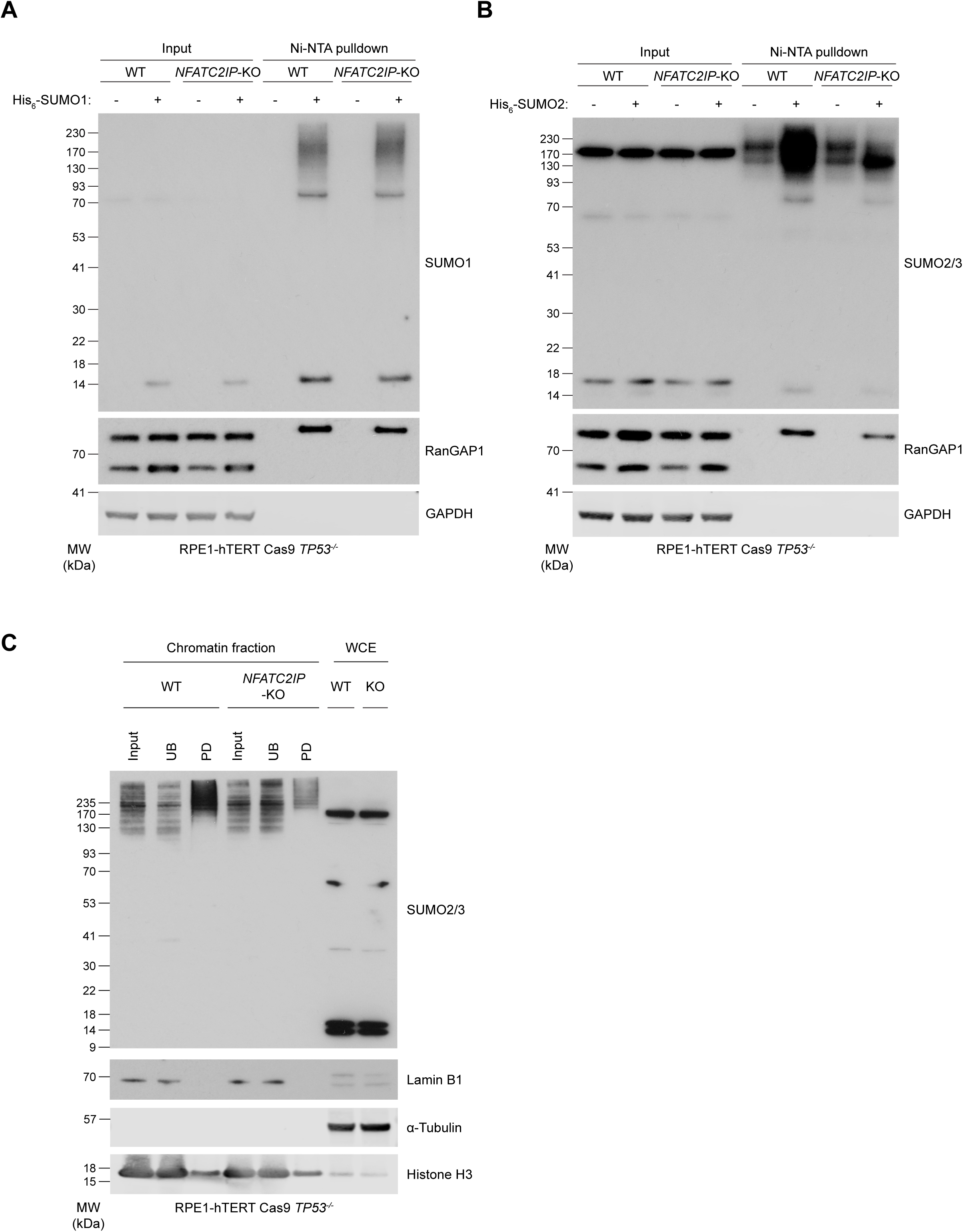
NFATC2IP promotes SUMOylation levels in chromatin. (A) and (B) Immunoblot analysis of whole-cell extracts of parental RPE1-hTERT Cas9 *TP53^-/-^*(WT) or isogenic *NFATC2IP*-KO cells that were transiently transfected with plasmids for overexpression of (A) His_6_-SUMO1 or (B) His_6_-SUMO2, followed by purification of His-tagged peptides using Ni-NTA agarose beads under denaturing conditions. The immunoblots were probed with antibodies to the indicated proteins. RanGAP1 was used as a control for SUMOylation. GAPDH was used as a loading control. (C) Immunoblot analysis of chromatin subfractions of extracts derived from parental RPE1-hTERT Cas9 *TP53^-/-^* (WT) or isogenic *NFATC2IP*-KO cells. SUMO-conjugated proteins were isolated by binding to the biotinylated S-Cap peptide, followed by affinity pulldown with streptavidin-conjugated magnetic beads. Input: input control fraction. UB: unbound supernatant after binding of biotin S-Cap to streptavidin beads. PD: proteins from S-Cap pulldown eluted from streptavidin beads. WCE: whole cell extract. Immunoblots were probed using antibodies to the indicated proteins. ɑ-tubulin, lamin B1, and histone H3 were included as controls for cytoplasmic, nucleoplasmic, and chromatin subcellular fractions, respectively.

Since the above experiments relied on overexpression of exogenous SUMO proteins, we next assessed whether NFATC2IP promotes SUMOylation under endogenous conditions. To do so, we subfractionated lysates of parental and *NFATC2IP*-KO cells and isolated SUMOylated proteins using a biotinylated S-Cap peptide that has high affinity for SUMOylated proteins. With this technique, we observed a large decrease in the amount of high molecular weight SUMO2/3-modified proteins specifically in the chromatin fraction of *NFATC2IP*-KO cells (Figure 7C) with no difference observed in the amount of SUMO2/3-modified proteins retrieved from either the nucleoplasmic or cytoplasmic fractions (Figure S8). Collectively, these data indicate that NFATC2IP promotes SUMOylation of chromatin-associated proteins, and based on the above interaction studies, that this activity may be related to its interaction with the SMC5/6 complex.

## Discussion

This work probed the genetic architecture of the response to compounds that perturb ubiquitin and Ubl conjugation pathways. This analysis identified a set 395 genes that modulate the fitness of human RPE1-hTERT *TP53^-/-^*cells to SUMO E1, ubiquitin E1, NEDD8 E1 and p97/VCP inhibition. As all four of these agents are or were investigated as anti-cancer agents in clinical trials, this dataset may offer new biomarkers of response or highlight the cellular pathways contributing to their anti-proliferative properties. In just one example, disruption of the *SPOP* gene, which is frequently mutated in prostate cancer (Zhang et al. 2023), sensitized RPE1 cells to the ubiquitin E1 inhibitor TAK-243 (Figure 1A).

The dataset can also be used to uncover new biological insights. As a first example, we validated the observation that TMED2 and TMED10, two transmembrane ER proteins, promote normal cellular resistance to p97/VCP inhibition. TMED2/10 are linked to various processes, including the retention of proteins such as Smoothened in the ER/Golgi (Di Minin et al. 2022), formation of lipid nanodomains (Anwar et al. 2022) and transport of GPI-anchored proteins (Bonnon et al. 2010). The exact mechanism by which loss of TMED2/10 causes a need for p97/VCP segregase activity is not clear, but we suspect this may be linked to the induction of an ER-associated unfolded protein response. In support of this possibility, mice with heterozygous mutation in *TMED2*, display a dilated ER and increased levels of eIF2α phosphorylation, which are indicative of ER stress (Hou et al. 2017). In the future it may be of interest to assess whether TMED2/TMED10 promotes ER protein quality control and whether this function is connected to p97 activity.

We focused the bulk of our validation work on defining the role of NFATC2IP in promoting the normal cellular resistance to SUMOylation inhibition by TAK-981. *NFATC2IP* disruption showed the largest effect size with respect to sensitization to TAK-981 in our cell line, outside disruption of the gene encoding the multidrug transporter MRP1. These results place NFATC2IP as a key mediator of the response of human cells to SUMO E1 inhibition.

While the developers of TAK-981 have highlighted its potential to stimulate anti-tumor immunity (Lightcap et al. 2021), we found that the intrinsic sensitivity of cells to SUMO E1 inhibition is largely driven by a few pathways that include genome maintenance and transcriptional regulation (Figure S3B). With respect to genome maintenance, in addition to NFATC2IP, our screen identified many genes encoding factors known to be involved in the resolution of recombination intermediates as promoting normal cellular resistance to SUMOylation inhibition, such as components of the BLM-RMI1-RMI2-TOP3A complex, GEN1 and the SMC5/6 complex. This is not entirely surprising given the key role of SUMO in controlling recombination in yeast (Ulrich et al. 2005; Branzei et al. 2006; Psakhye and Jentsch 2012). Indeed, our data is consistent with a model where NFATC2IP promotes SUMOylation of one or more proteins involved in resolving either recombination intermediates or topological entanglements during interphase. In support of this model, we find that NFATC2IP loss exacerbates the genome instability phenotypes seen with TAK-981 treatment, and we observed that NFATC2IP promotes SUMOylation of chromatin-associated proteins. Although we have not been able to identify proteins whose SUMOylation status is specifically influenced by NFATC2IP, we anticipate that a subset of them will be substrates of NSMCE2, the SMC5/6-associated SUMO E3 ligase. Indeed, SMC5 interacts with NFATC2IP, with the NFATC2IP SLD2 modeled to be in a prime position to assist with NSMCE2-dependent SUMOylation by either positioning the SUMO E2 UBC9 or by promoting NSMCE2 catalytic activity in a manner similar to the described backside SUMO interaction with UBC9 (Varejao et al. 2021) (Figure 6E). Interestingly, in addition to the SLD2, we identified another region essential for NFATC2IP action located within residues 101-140. The molecular function of this region remains unclear as it was not modeled to interact any other member of the SMC5/6 pathway. However, we note that NFATC2IP homologs have been shown to display DNA-binding activity that maps N-terminal of the SLD1/2 domains (Urulangodi et al. 2015; Sebesta et al. 2017). Whether this region in NFATC2IP also confers DNA-binding activity is currently unknown but represents an attractive starting point for future studies.

Finally, given that the phenotypes of *NFATC2IP-*KO cells treated with low doses of the SUMO E1 inhibitor are remarkably similar to the phenotypes associated with mutations in SMC5/6 or associated proteins (Gallego-Paez et al. 2014; Payne et al. 2014; Jacome et al. 2015; Pryzhkova and Jordan 2016), it may be of interest to screen for *NFATC2IP* mutations in patients that display chromosome breakage disorders linked to the SMC5/6 complex such as Atelis, Seckel and LICS syndromes, associated with *SMC5*, *SLF2*, *NSMCE2* and *NSMCE*, respectively. However, as the *NFATC2IP-KO* cellular phenotypes are only uncovered when SUMOylation is perturbed, we expect that the phenotypes associated with loss of NFATC2IP will be milder than those seen with mutations in SMC5/6 complex-coding genes or will affect tissues that have lower levels of SUMOylation than our model cell line.

## Acknowledgements

We thank Rachel Szilard for critical reading of this manuscript and other members of the Durocher lab for valuable discussion. We thank Oscar Fernandez-Capetillo for alerting us to the similarity between our phenotypes and the chromatin bridges observed in *NSMCE2*-deficient cells, and Niels Mailand for sharing results prior to their publication. D.D. is a Canada Research Chair (Tier 1) and work in the D.D. lab was funded from grants from the Canadian Institutes for Health Research (CIHR, PJT-180438) and the Reiss Institute for Healthy Aging Research.

## Author contributions

Conceptualization, T.C., M.O. and D.D.; Investigation, T.C., Y.Z. M.O. and D.S.; Writing – Original Draft, T.C. and D.D.; Writing – Review and Editing, T.C. Y.Z. and D.D.; Visualization, T.C., Y.Z., L.H., and D.S.; Formal Analysis, L.H. and D.S; Data curation, L.H.; Supervision, D.D.; Funding Acquisition, D.D.

## Supplementary Figure Legends

**Figure S1.**
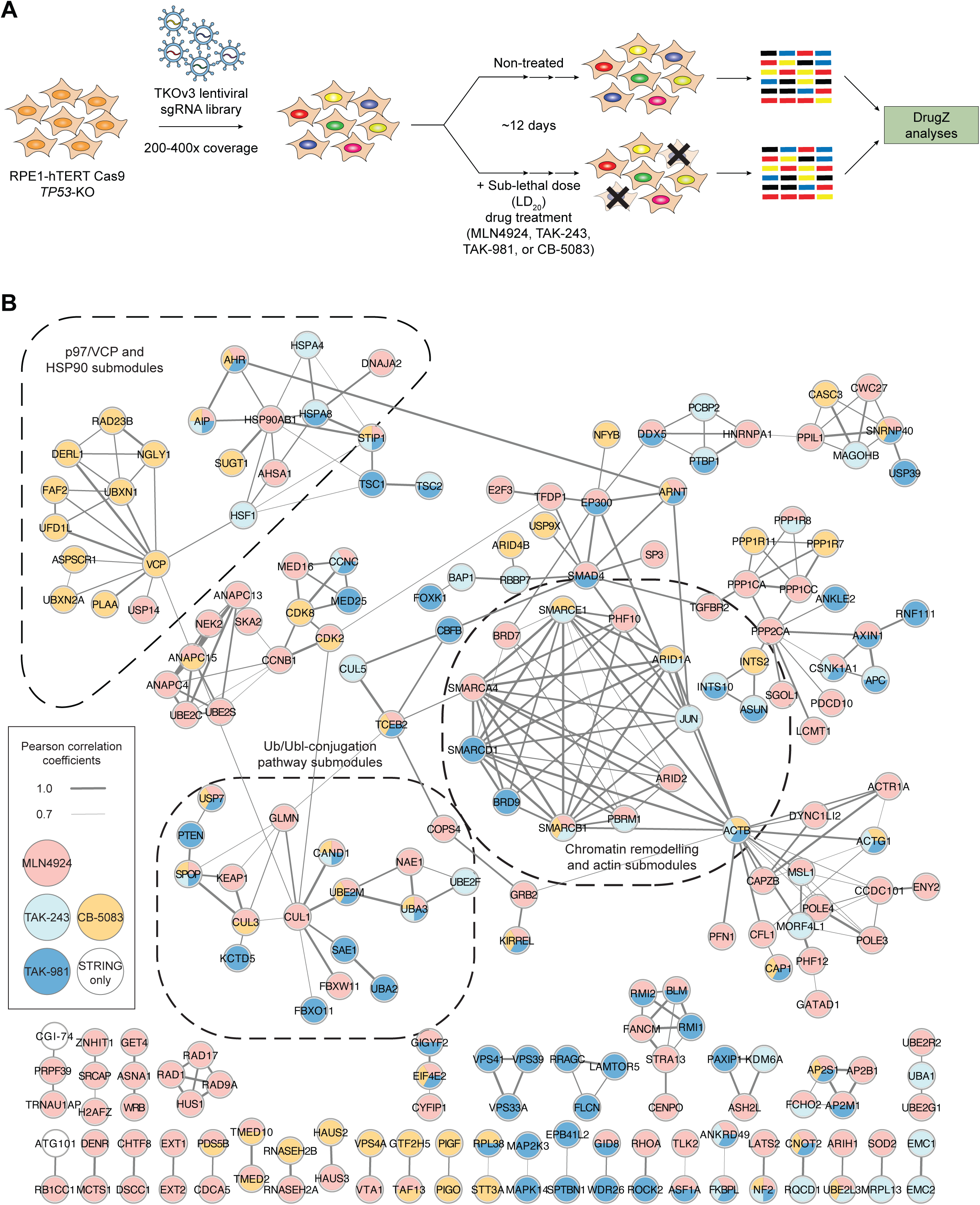
Protein-protein interaction network of the screen hits. (A) Schematic of the chemogenomic CRISPR screen performed in RPE1-hTERT Cas9 *TP53^-/-^* cells. (B) Network of genes who products have known protein-protein interactions among the 395 hit genes identified in the chemogenomic CRISPR screens with Ub/Ubl pathway inhibitor at a Pearson correlation coefficient (PCC) value greater than 0.7. A total of 210 genes were included in the network based on their protein products having experimentally examined and database-annotated protein associations in the STRING database (version 4.0) with the protein products of other genes in the hit gene set. Colours of nodes represent specific chemogenomic CRISPR screen(s) in which a particular gene has been identified as a hit. Size of edge connecting different nodes indicates correlation level between nodes. Three clusters of interconnected submodules are highlighted with their known functions.

**Figure S2.**
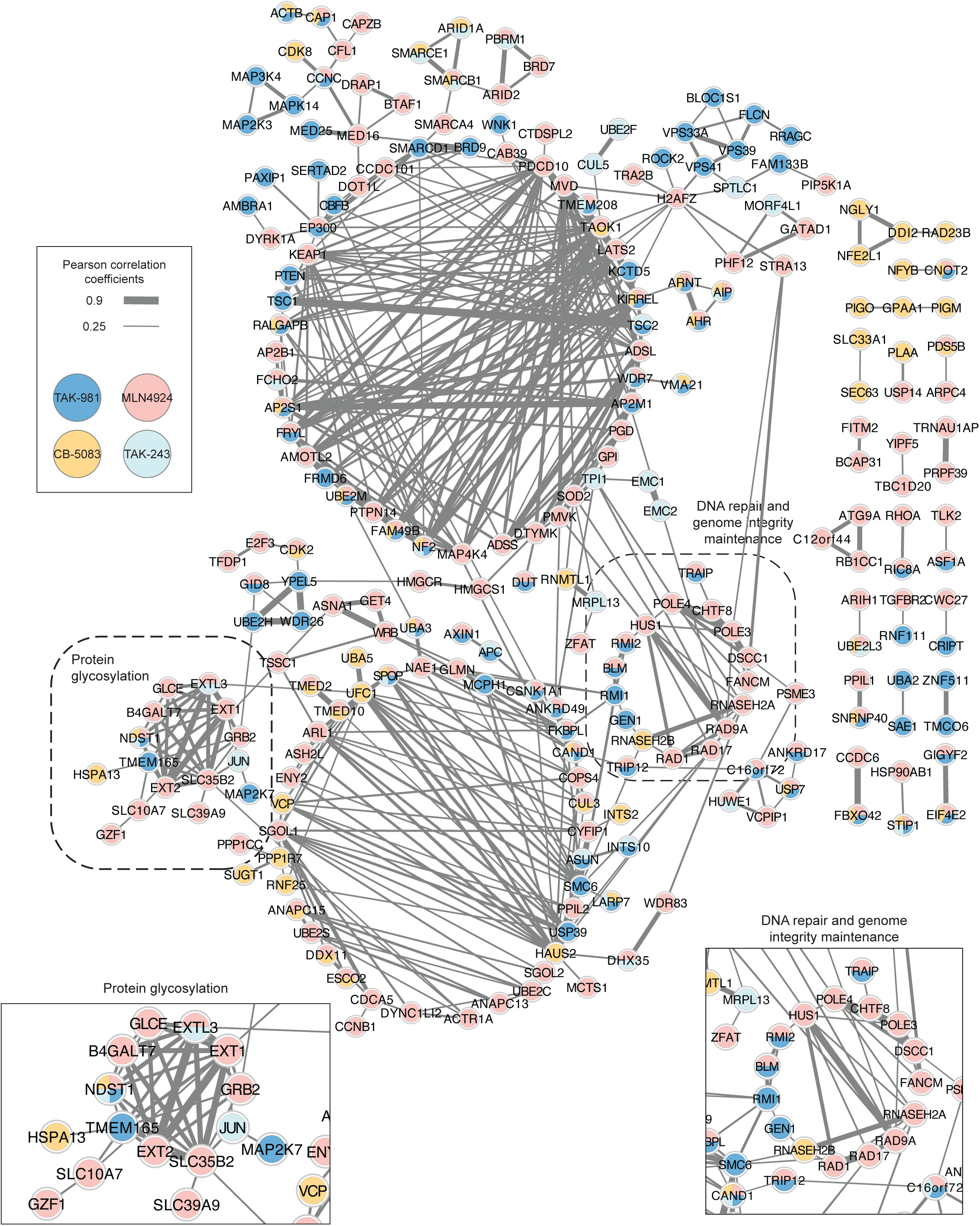
Gene-gene essentiality correlation network of the screen hits. (A) Network of 240 genes, among the 395 hit gene set from the Ub/Ubl inhibitor CRISPR screens, whose gene essentiality scores are correlated with another gene at a PCC value greater than 0.25 based on the DepMap database version 22Q4. Colours of nodes indicate specific inhibitor CRISPR screen(s) in which a particular gene has been identified as a hit. Size of edge between different nodes represents the level of correlation between two nodes. Two highlighted clusters are enriched with genes whose protein product is involved in protein glycosylation or DNA damage responses.

**Figure S3.**
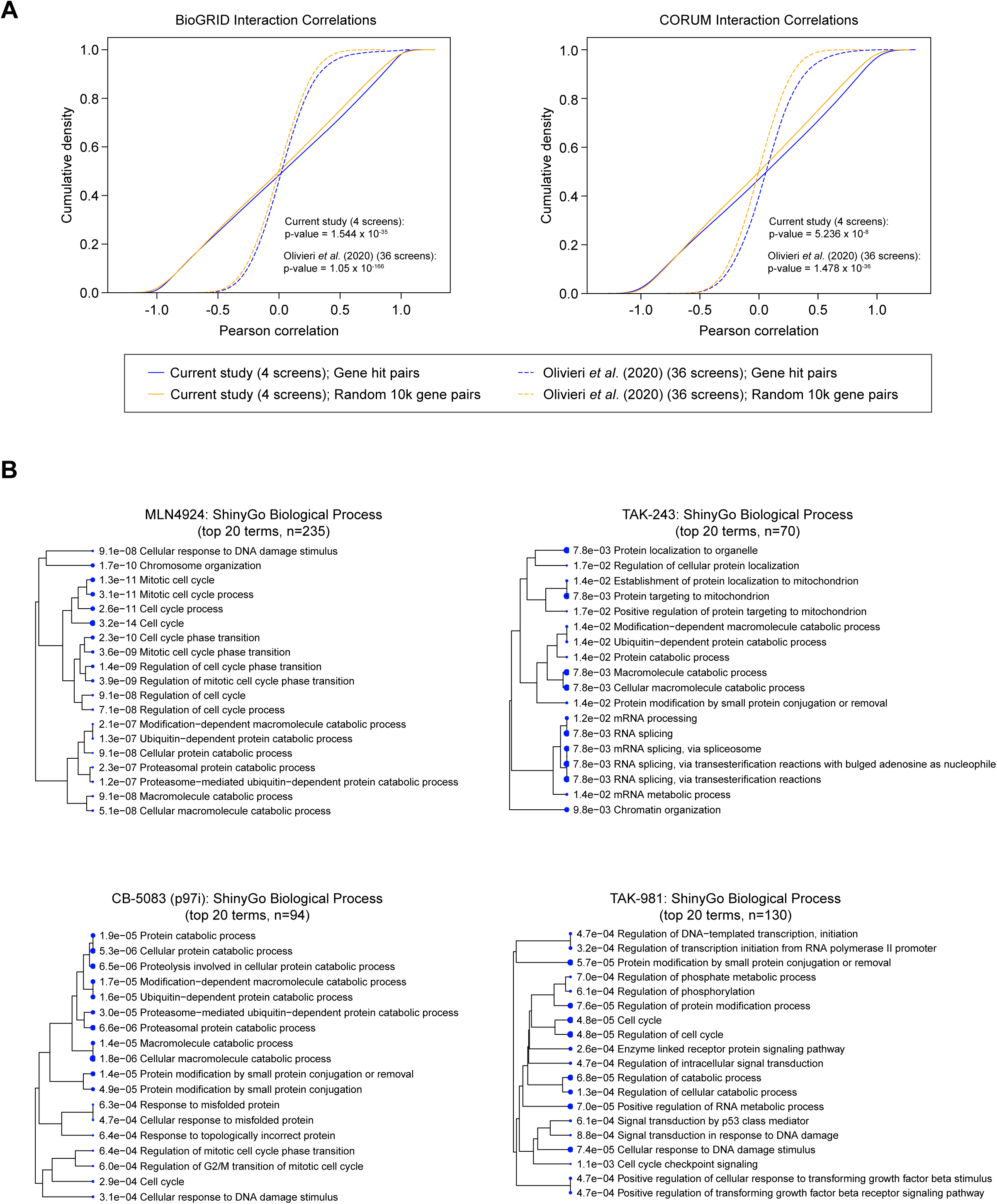
Gene ontology enrichment analyses. (A) Enrichment of genes among the 395 hit gene set whose gene products are annotated for protein interactions with other genes using either BioGRID version 4.4.218 (left) or CORUM version 4.1 (right) protein interaction database. Pairs of genes were taken from the indicated protein interaction databases, and gene pairs that have at least one gene identified in our 395 hit gene set were assigned as ‘Gene hit pairs’. 10,000 gene pairs that did not include a gene present in our list of hit genes were assigned as ‘Random 10k gene pairs’. For each gene pair set, the Pearson correlation was calculated between normZ of the two genes in each gene pair and plotted as cumulative density functions. For statistical testing of the difference between the two distributions of hit gene pairs and random gene pairs, the Kolmogorov-Smirnov test was used to calculate the p-values. The results using the 890 hit gene set in Olivieri et al. (2020) are plotted (dashed lines) with the associated *P*-values. (B) Analysis of gene ontology enrichment of biological process terms using ShinyGo among the hit genes identified in the indicated Ub/Ubl inhibitor CRISPR screens (Ge et al. 2020). An FDR cut-off of 0.05 was used, and the top 20 enriched functional terms are shown with their individual *P-*values. The relative size of the circle for each GO term represents significance of *P-* value.

**Figure S4.**
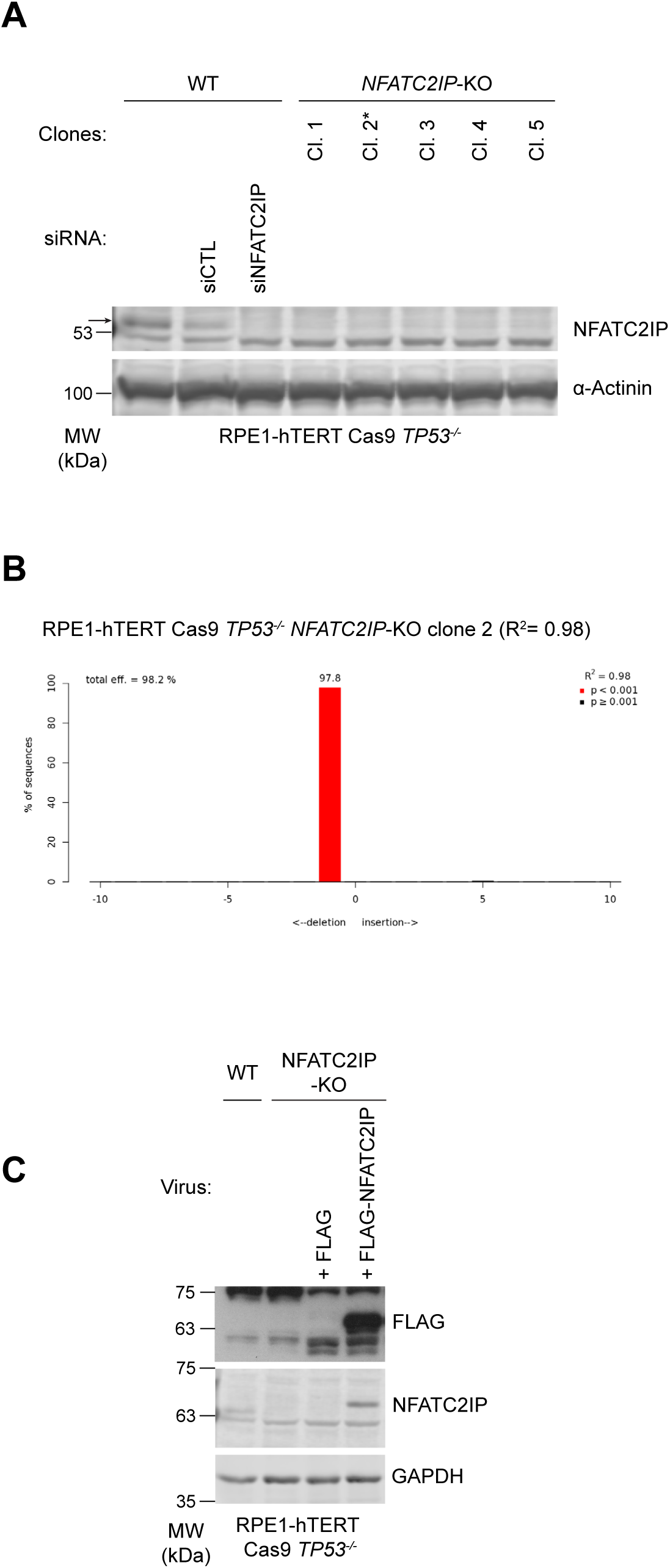
Validation of the *NFATC2IP* clonal KO. (A) Immunoblot analysis of the clonal *NFATC2IP*-KO cell lines isolated from parental RPE1-hTERT Cas9 *TP53^-/-^* (WT) cells after transfection of a Cas9 ribonucleoprotein complex assembled with the *NFATC2IP*-targeting sgRNA2 (Table S4). Arrow indicates the endogenous NFATC2IP protein band. Cells transfected with an siRNA targeting *NFATC2IP* mRNA along with a non-targeting siRNA (siCTL) were included to validate the NFATC2IP antibody. ɑ-Actinin was used as a loading control. Asterisk (*) indicates the *NFATC2IP*-KO clone of RPE1-hTERT Cas9 *TP53^-/-^* cells that was used for all further experiments. (B) Genotyping of the *NFATC2IP*-KO clone 2 of RPE1-hTERT Cas9 *TP53^-/-^* cell line using Tracking of Indels by Decomposition (TIDE) analysis (Brinkman et al. 2014). Bars represent the percentage of the sequence of the PCR product amplified around the CRISPR-targeted genomic locus that contains a particular indel mutation. The red colour of the major bar indicates that the *P*-value is less than 0.001, with a R^2^ value of 0.98. (C) Immunoblot analysis to validate expression of epitope-tagged NFATC2IP protein. Whole cell lysates from parental RPE1-hTERT Cas9 *TP53^-/-^* (WT) or isogenic *NFATC2IP*-KO cells that were left untransduced or were transduced with a lentivirus encoding Flag only or Flag-tagged full-length NFATC2IP protein were analyzed by immunoblotting with antibodies to FLAG, NFATC2IP, or GAPDH (loading control).

**Figure S5.**
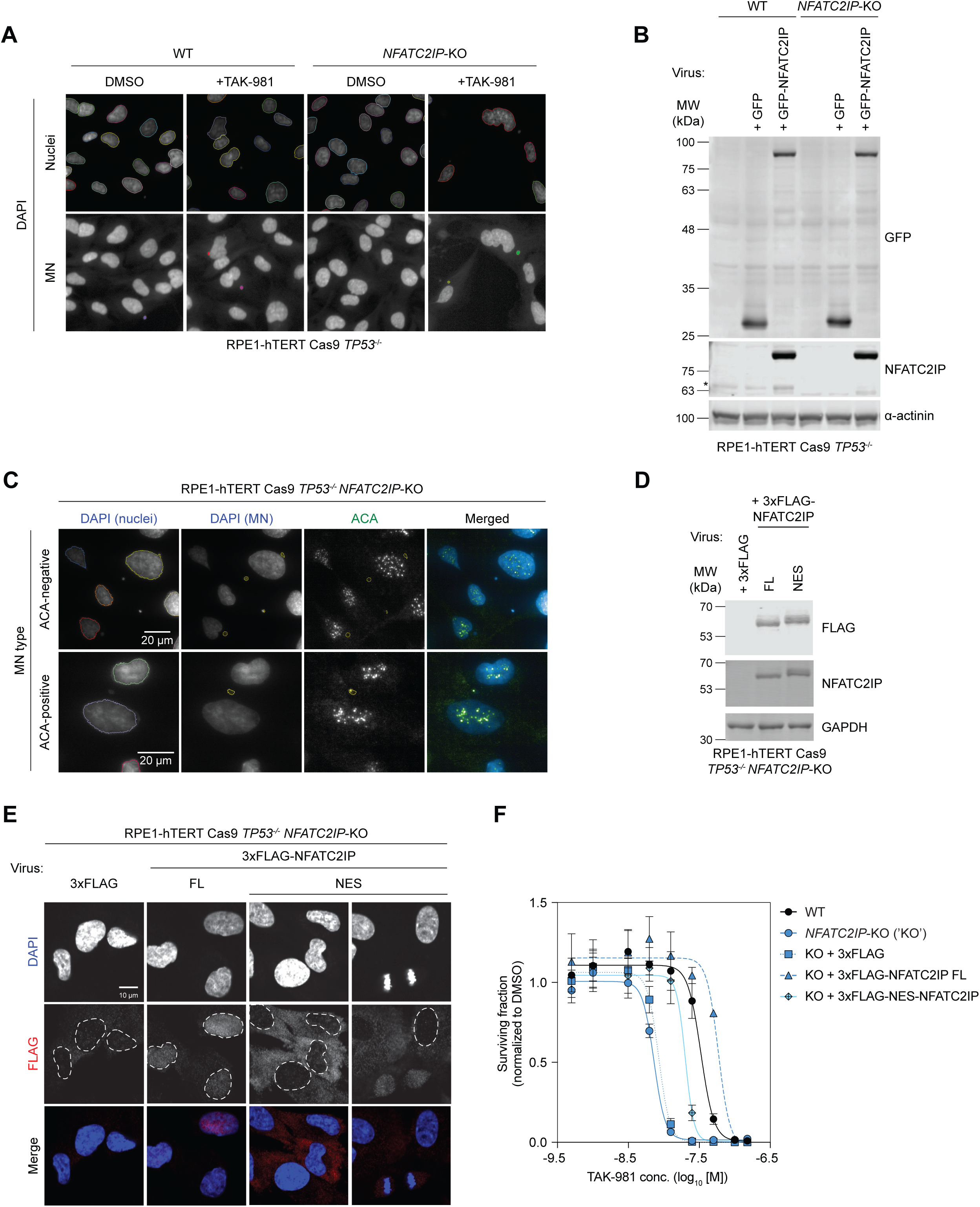
NFATC2IP functions in interphase to suppress genome instability upon SUMOylation inhibition. (A) Representative micrographs from automated quantitation of MN formation after treatment with increasing doses of TAK-981 for 72 h, related to Figure 4A. The DAPI-stained parental RPE1-hTERT Cas9 *TP53^-/-^* (WT) or isogenic *NFATC2IP*-KO cells were imaged using an InCell 6000 Analyzer with a 20x objective. The resulting images were used for generating initial nuclear masks (outlined in ‘nuclei’ image panels) and MN masks (outlined in ‘MN’ image panels) using the Columbus image analysis suite. (B) Immunoblot analysis to validate expression of exogenous NFATC2IP protein; related to Figure 4A. Whole cell lysates from parental RPE1-hTERT Cas9 *TP53^-/-^* (WT) or isogenic *NFATC2IP*-KO cells that were left untransduced or transduced with lentiviruses encoding GFP only or GFP-tagged full-length NFATC2IP protein were immunoblotted with antibodies to GFP, NFATC2IP, or ɑ-actinin (loading control). Asterisk indicates endogenous NFATC2IP. (C) Representative immunofluorescence micrographs from automated quantitation of MN formation and MN type determination after treatment with TAK-981 (25 nM) for 48 h; related to Figure 4B. RPE1-hTERT Cas9 *TP53^-/-^* (WT) or isogenic *NFATC2IP*-KO cells were stained with DAPI and anti-centromeric antibody (ACA). Images were acquired from DAPI and FITC (ACA) channels using an InCell 6000 Analyzer with a 60x objective. Images from the DAPI channel were used for generating initial nuclear masks (outlined in ‘DAPI (nuclei)’ panels) and the MN mask (outlined in ‘DAPI (MN)’ panels). Following identification of MN, the number of MN that are positive or negative for ACA signal were quantitated using the Columbus image analysis suite from the FITC (ACA) channel. Scale bars=20 µm. (D) Immunoblot analysis to validate expression of exogenous NFATC2IP proteins; related to Figure 4F. Whole cell lysates from RPE1-hTERT Cas9 *TP53^-/-^ NFATC2IP*-KO cells that were transduced with lentiviruses encoding 3xFlag only or 3xFlag-tagged full-length or NES-tagged NFATC2IP protein were immunoblotted with antibodies to FLAG, NFATC2IP, or GAPDH (loading control). FL: full-length NFATC2IP. NES: NES-NFATC2IP. (E) Representative micrographs of immunofluorescence analysis to examine localization of exogenous Flag-tagged NFATC2IP proteins; related to Figure 4F. RPE1-hTERT Cas9 *TP53*^-/-^ *NFATC2IP*-KO cells were stained with the indicated antibodies and DAPI as a nuclear marker. FL: full-length NFATC2IP. NES: NES-NFATC2IP. Scale bar=10 µm. (F) Surviving fractions (normalized to the DMSO control condition) from TAK-981 dose-response clonogenic survival assays in parental RPE1-hTERT Cas9 *TP53^-/-^* (WT) or isogenic *NFATC2IP*-KO cells that were left untransduced or were transduced with lentiviruses encoding exogenous full-length (FL) or NES-fused NFATC2IP protein. Related to Figure 4F. Data are shown as the mean ± s.e.m. (n=3).

**Figure S6.**
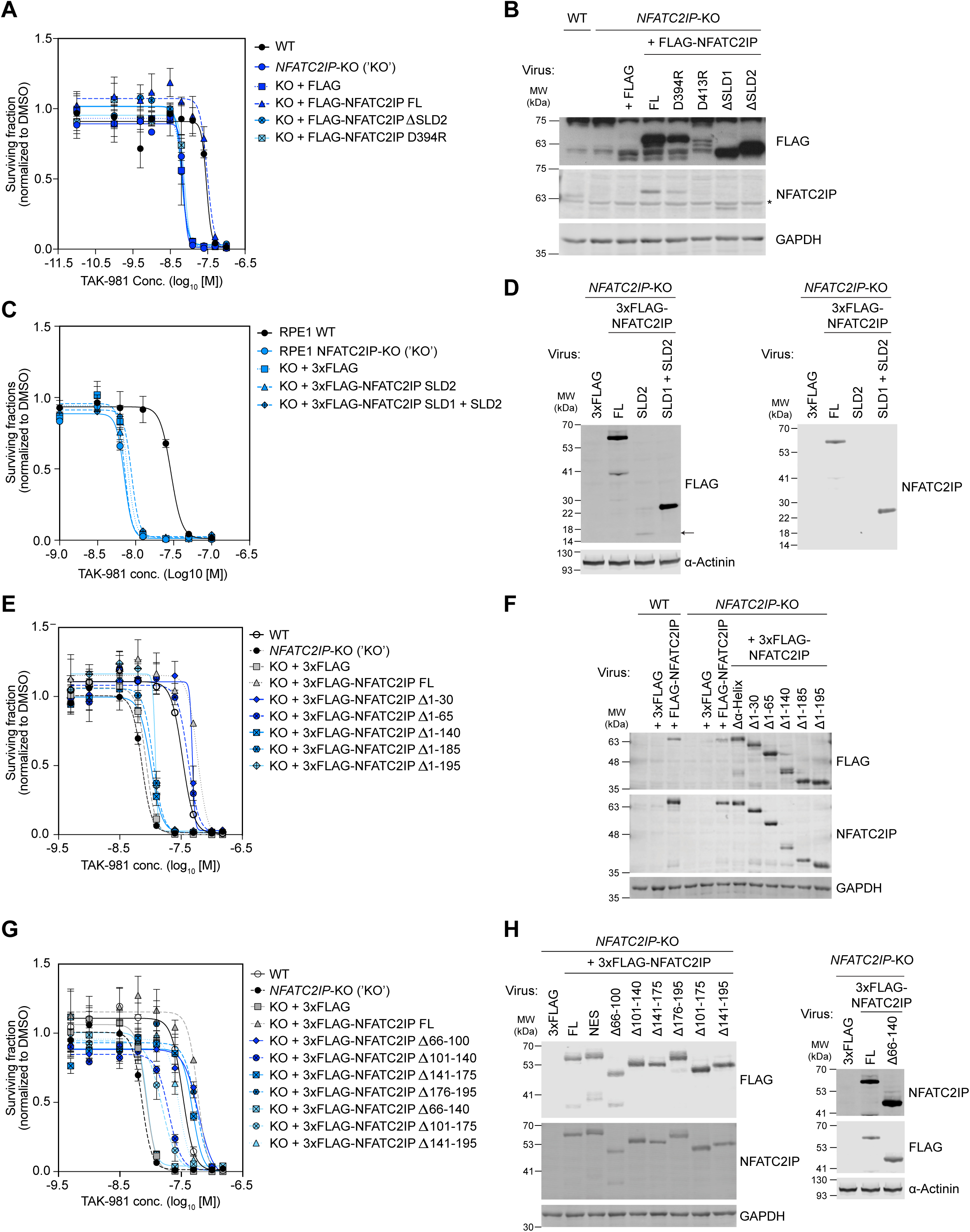
Supporting data for the structure-function analysis of NFATC2IP. Results of TAK-981 dose-response clonogenic survival assays (panels (A), (C), (E), and (G)) accompanied by immunoblot analysis of whole cell lysates (panels (B), (D), (F), and (H)) to validate the expression of exogenous NFATC2IP constructs. Parental RPE1-hTERT Cas9 *TP53^-/-^*(WT) or isogenic *NFATC2IP*-KO cells were left untransduced or were transduced with the indicated constructs. Data in plots of surviving fractions are normalized to the DMSO (vehicle) condition and are shown as the mean ± s.e.m. (n=3). Immunoblots were probed using antibodies to FLAG orNFATC2IP, along with GAPDH or ɑ-actinin as loading controls. The NFATC2IP antibody recognizes SLD2 of the NFATC2IP protein. FL: full-length NFATC2IP. NES: NES-NFATC2IP. (A) –(D) Analysis of the SLD1 and SLD2 of NFATC2IP; related to Figure 5A–C. Asterisk beside NFATC2IP immunoblot in (B) indicates non-specific bands. Arrow in (D) indicates the migration of the 3xFLAG-NFATC2IP SLD2 protein. (E) –(H) Analysis of NFATC2IP proteins with the indicated deletion mutations schematized in Figure S7; related to Figure 5D.

**Figure S7.**
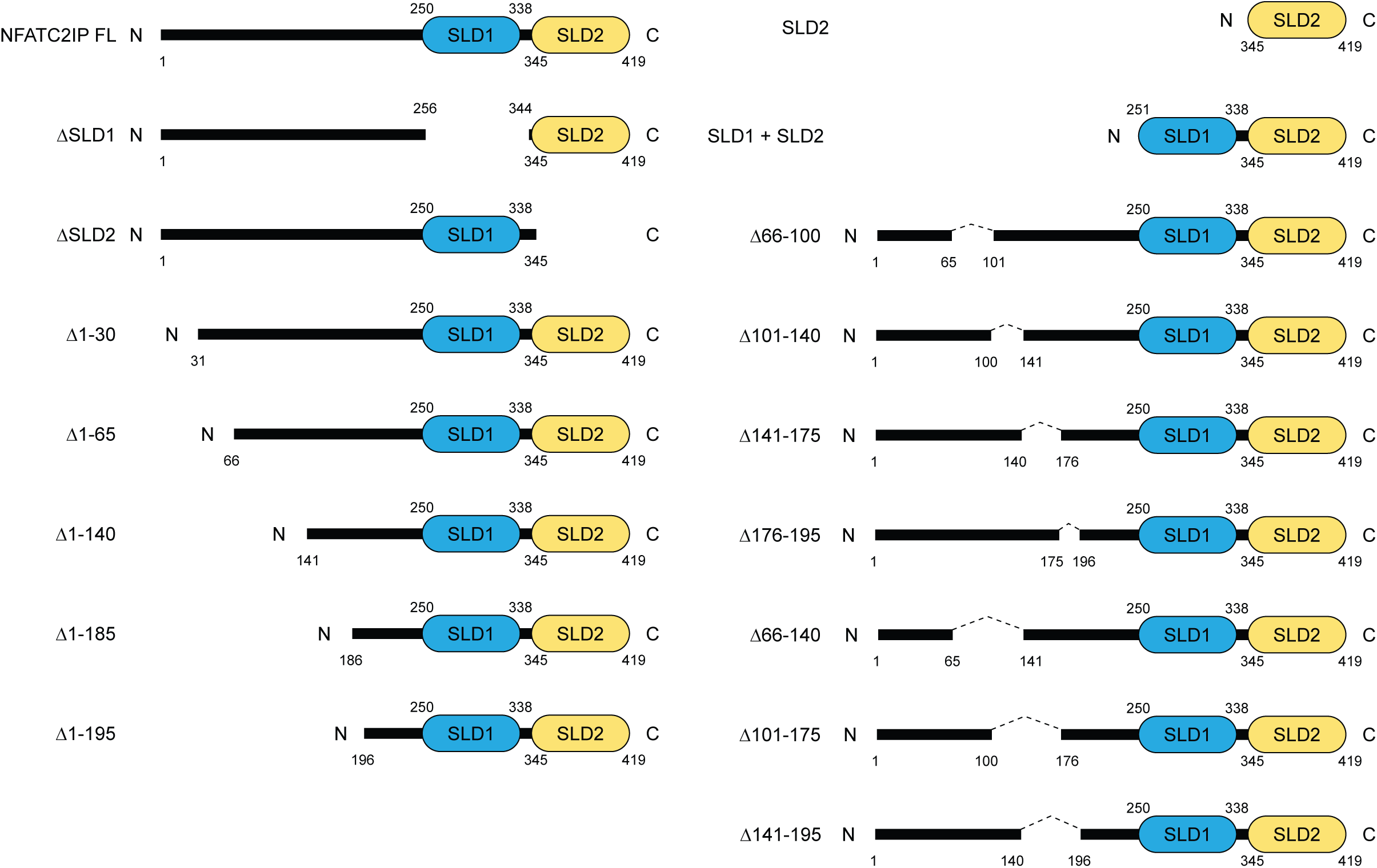
Schematics of the NFATC2IP deletion mutants. Schematic summary of the NFATC2IP proteins with various deletion mutations that were used in the experiments to elucidate functions of NFATC2IP SLDs and to identify regions outside of SLD1 – SLD2 that contribute to the response of NFATC2IP to SUMOylation pathway inhibition.

**Figure S8.**
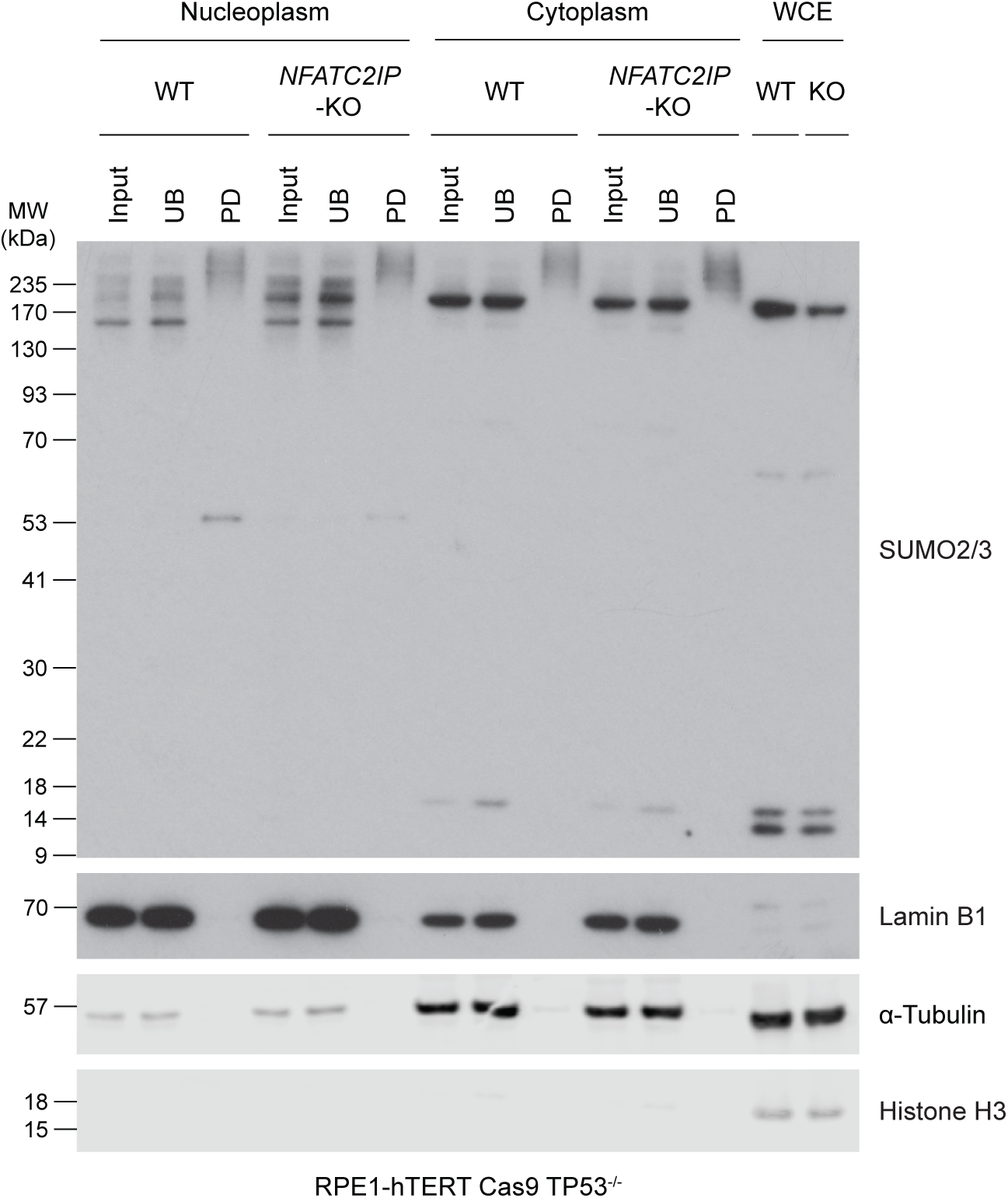
Analysis of SUMOylation levels of nucleoplasmic and cytoplasmic proteins, related to Figure 7C. Immunoblot analysis of nucleoplasmic and cytoplasmic subfractions of extracts derived from parental RPE1-hTERT Cas9 *TP53*^-/-^ (WT) or isogenic *NFATC2IP*-KO cells. SUMO-conjugated proteins were isolated by binding to the biotinylated S-Cap peptide, followed by affinity pulldown using streptavidin-conjugated magnetic beads. Input: input control fraction. UB: unbound supernatant after binding of biotin S-Cap to streptavidin beads. PD: proteins from S-Cap pulldown eluted from streptavidin beads. WCE: whole cell extracts. Immunoblots were probed with antibodies to the indicated proteins. ɑ-Tubulin, Lamin B1, or Histone H3 were included as controls for cytoplasmic, nucleoplasmic, or chromatin subcellular fractions, respectively.

## Methods

### Cell lines and cell culture

RPE1-hTERT Cas9 *TP53-*KO and 293T cell lines were cultured in Dulbecco’s Modified Eagle Medium (DMEM, Gibco, cat # 11965092) supplemented with 10% (v/v) FBS (Wisent Bioproducts, cat # 080-150) and 1% (v/v) penicillin/streptomycin (Wisent Bioproducts, cat # 450-201-EL) and grown at 37℃ and 5% CO_2_. Where indicated, carfilzomib (Selleckchem, cat # S2853) was added at a concentration of 20 nM for 4 h. The clonal RPE1-hTERT Cas9 *TP53^-/-^ NFATC2IP*-KO cell lines were generated by transfecting a ribonucleoprotein (RNP) complex of the *NFATC2IP*-targeting sgRNA2 (Table S4) and purified Cas9 protein into RPE1-hTERT Cas9 *TP53^-/-^* cells using Lipofectamine CRISPRMAX Cas9 Transfection Reagent (Invitrogen, cat # CMAX00003). Transfection of the RNP complex was performed according to the manufacturer’s protocol for 24 h and seeded for clonal isolations. The selected *NFATC2IP*-KO clones were validated for successful gene editing by TIDE analysis (Brinkman et al. 2014) and by probing for endogenous NFATC2IP protein levels by immunoblotting.

### Plasmids

For CRISPR-mediated gene perturbation, DNA oligonucleotides containing sgRNAs were cloned into LentiCRISPRv2 (Addgene # 52961) using BsmBI. For generating NFATC2IP-expressing plasmids, a plasmid containing the human *NFATC2IP* cDNA was purchased from Lunenfeld-Tanenbaum Research Institute OpenFreezer repository (OpenFreezer ID V84563). The *NFATC2IP* coding sequence was amplified by PCR with primers containing flanking AscI and EcoRV restriction sites and cloned into pcDNA5-FRT/TO-FLAG, pcDNA5-FRT/TO-eGFP, and pcDNA5-FRT/TO-3xFLAG for N-terminal epitope tagging. To generate plasmids expressing NFATC2IP mutants, site-directed mutagenesis was performed via PCR. The sequences encoding FLAG- or eGFP-tagged NFATC2IP were then PCR-amplified with primers containing flanking NotI and XbaI restriction sites and cloned into the pHIV-NAT-T2A-hCD52 lentiviral vector (kind gift from Dr. R. Scully). For generating NSMCE2-expressing plasmids, a plasmid with the human *NSMCE2* cDNA was purchased from GenScript (cat # Ohu31586D). The *NSMCE2* coding sequence was PCR-amplified to add a Kozak consensus sequence for efficient initiation of translation and flanking KpnI and XhoI restriction sites, and then cloned into pcDNA3.1-C-2xHA vector (kind gift from Dr. R McInnes) for tagging the epitope at its C-terminal end.

### Lentiviral transduction

To produce lentivirus, 4 × 10^6^ 293T cells were seeded in a 10-cm plate 1 d prior to transfection and co-tranfected with the 3^rd^ generation lentiviral packaging plasmids (5 µg pVSV.G, 3 µg pMDLg/pRRE, and 2.5 µg pRSV-Rev; Addgene # 14888, 12251, and 12253, respectively) plus 10 µg of the vector of interest using Mirus TransIT-LT1 transfection reagent (Mirus Bio LLC, cat # MIR 2305). Medium was refreshed 16 h post-transfection, and the viral supernatant was collected approximately 36-40 h post-transfection and passed through a 0.45-µm filter. For infection of target cells, the viral supernatant was supplemented with 8 µg/ml polybrene (Sigma, cat # H9268). For selection of transduced RPE1-hTERT Cas9 *TP53^-/-^* WT or *NFATC2IP*-KO cells, cells were grown in media containing 1.5 µg/ml of puromycin (Life Technologies, cat # A1113802) for 48 h, or 400 µg/ml of nourseothricin (NAT, Gold Biotech, cat # N-500) for 4–5 days.

### CRISPR Cas9 screens

Chemogenomic CRISPR screens were performed by following the protocol described in (Olivieri and Durocher 2021). Briefly, RPE1-hTERT Cas9 *TP53^-/-^* cells were transduced with the TKOv3 lentiviral sgRNA library (Hart et al. 2017) at a multiplicity of infection (MOI) of ∼ 0.3, and selected with puromycin (1.5 μg/ml) for 48 h. The pooled population of transduced cells were subcultured every 3 d in two technical replicates until 6 d post-selection (t6). At the t6 timepoint, the subpopulations of cells were either exposed to DMSO (vehicle) as a non-treated control or treated with the Ub/Ubl pathway inhibitors at the dosages described in Table S1 for 12 d. Cells were harvested at the t18 timepoint, genomic DNA was purified using the QIAmp DNA Blood Maxi kit (QIAGEN, cat # 51194) and the integrated sgRNA sequences were amplified and barcoded by two-step PCR using NEBNext Ultra II Q5 Master Mix (New England Biolabs, cat # M5044). The barcoded sgRNA samples were sequenced on an Illumina NextSeq500 to quantitate representation of each sgRNA in the Ub/Ubl pathway inhibitor-treated or non-treated control samples. As described previously, the gene-level normZ scores were computed using DrugZ from the readcounts (Table S5) (Colic et al. 2019).

### Clonogenic survival assays

For clonogenic survival assays with TAK-981, cells were seeded in media containing a range of TAK-981 concentrations at the seeding density of 400 cells per 10-cm plates. TAK-981-containing media was refreshed every 4 d, and cells were grown for 14 d. Then, cells were rinsed with DPBS (Gibco, cat # 14190144) and stained with 0.5% (w/v) crystal violet/20% methanol for 30 min. Numbers of colonies were counted by images analysis using a GelCount (Oxford Optronix) and the relative surviving fractions were plotted by normalizing to DMSO controls.

For clonogenic survival assays with CB-5083, cells were lentivirally transduced with an sgRNA-expression construct and selected with puromycin (1.5 μg/ml) for 48 h. 72 h post-selection, the transduced cells were seeded in media containing 200 nM CB-5083 at a seeding density of 500 cells per 10-cm plate and were grown for 13 d. CB-5083-containing media was refreshed every 6 days. Colonies were stained and quantitated as described above.

### Quantitative reverse transcription (qRT) PCR

RNA was isolated from RPE1-hTERT Cas9 *TP53^-/-^* cells using an RNeasy kit (Qiagen, cat # 74104), and cDNA was synthesized using SuperScript™ III Reverse Transcriptase (Invitrogen, cat # 18080093). The prepared cDNA was used as a template for qRT-PCR that was performed using TaqMan Gene Expression Assays (Thermo). The following TaqMan probes were used in the experiment (detailed sequence information in Table S4): PSMA7 (Hs00895424_m1), PSMB7 (Hs00160607_m1), PSMC4 (Hs00197826_m1), and GAPDH (Hs99999905_m1).

### Antibodies

Primary antibodies used in this study were the following: rabbit anti-NRF1 (Cell Signaling Technologies, D5B10, cat # 8052, 1:1000), rabbit anti-GAPDH (Sigma, cat # G9545, 1:10000), mouse anti-NFATC2IP (Santa Cruz Biotechnology, B-1, cat # sc-377461, 1:100), rabbit anti-GFP (Abcam, cat # ab290, 1:1000), mouse anti-ɑ-tubulin (Cell Signaling Technologies, DM1A, cat # 3873, 1:2000), anti-centromere protein antibody (ACA) (Antibodies Incorporated, cat # 15-235, 1:1000), rat anti-FLAG (BioLegend, L5, cat # 637301, 1:500 for immunofluorescence), mouse anti-FLAG M2 (Sigma, cat # F1804, 1:500 for immunoblotting), rabbit anti-NSMCE2 (Novus Biologicals, cat # NBP1-76263, 1:1000), rabbit anti-SMC5 (Novus Biologicals, cat # NB100-469, 1:500), rabbit anti-RanGAP1 (Abcam, EPR3295, cat # ab92360, 1:2000), rabbit anti-SUMO1 (Abcam, Y299, cat # ab32058, 1:1000), rabbit anti-SUMO2/3 (Abcam, cat # ab3742, 1:1000), rabbit anti-Lamin B1 (Abcam, cat # ab16048, 1:1000), and rabbit anti-Histone H3 (Abcam, cat # ab1791, 1:4000).

For immunoblotting, the following secondary antibodies were used: horseradish peroxidase (HRP)-conjugated AffiniPure goat anti-mouse IgG (H+L) (Jackson ImmunoResearch Laboratories Inc., cat # 115-035-003, 1:5000), HRP-conjugated AffiniPure goat anti-rabbit IgG (H+L) (Jackson ImmunoResearch Laboratories Inc., cat # 111-035-144, 1:5000), IRDye 680RD donkey anti-mouse IgG (LI-COR Biosciences, cat # 926-68072, 1:5000), IRDye 680RD goat anti-mouse IgG (LI-COR Biosciences, cat # 926-68070, 1:5000), and IRDye 800CW donkey anti-rabbit IgG (LI-COR Biosciences, cat # 926-32213, 1:5000). For immunofluorescence, the secondary antibodies used in this study include the following: Alexa Fluor 555 goat anti-rabbit IgG (H+L) (Invitrogen, cat # A21428, 1:1000), Alexa Fluor 555 goat anti-rat IgG (H+L) (Invitrogen, cat # A21434, 1:1000), Alexa Fluor 647 goat anti-rabbit IgG (H+L) (Invitrogen, cat # A21244, 1:1000), Alexa Fluor 647 goat anti-mouse IgG (Invitrogen, cat # A21236, 1:1000), and Alexa Fluor 488 goat anti-human IgG (H+L) (Invitrogen, cat # A11013, 1:1000).

### Immunofluorescence

Cells were seeded on glass coverslips and subjected to various treatments detailed elsewhere. Cells on glass coverslips were washed with PBS and fixed with 4% (w/v) formaldehyde/PBS (Pierce, 16% formaldehyde (w/v), methanol-free, Thermo Fisher, cat # 28908, diluted in PBS) for 15 min and then permeabilized with 0.3% (v/v) Triton X-100/PBS for 30 min. Cells were incubated with PBG blocking buffer (0.2% fish gelatin and 0.5% (w/v) bovine serum albumin (BSA), diluted in PBS) for 30 min, and subsequently with primary antibodies diluted in PBG blocking buffer for 1-2 h. Following three washes with PBS, cells were incubated with PBG blocking buffer containing secondary antibodies and 0.4 μg/ml DAPI (4’,6-diaminido-2-phenylindole, Sigma, cat # D9542) for 1 h, and subsequently washed with PBS three times. Cells were mounted onto glass slides using ProLong Gold Antifade mountant (Thermo Fisher, cat # P36930) and imaged on a Zeiss LSM780 confocal microscope.

### Quantitation of micronuclei formation

Cells were seeded in 96-well plates at a seeding density of 3000 cells per well, 24 h prior to the TAK-981 treatment. Cells were treated with TAK-981 for 48 h, followed by fixation with 4% (w/v) formaldehyde/PBS (Pierce, 16% formaldehyde (w/v), methanol-free, Thermo Fisher, cat # 28908, diluted in PBS) for 15 min and permeabilization with 0.3% (v/v) Triton X-100/PBS for 30 min. Cells were rinsed with PBS and incubated for 30 min with blocking buffer containing 5% (w/v) BSA and 0.1% (v/v) Tween-20 diluted in PBS. Cells were incubated with primary antibodies diluted in blocking buffer for 2 h, followed by three washes with PBT (0.1% (v/v) Tween-20/1xPBS). Subsequently cells were incubated with the blocking buffer containing secondary antibodies and DAPI. After final washing with PBT three times and with PBS once, 200 μl PBS was added to each well. Plates were scanned for image acquisitions on an InCell Analyzer 6000 automated microscope (GE Healthcare) with 20x or 60x objectives. Image analyses for micronuclei quantitation were performed using Columbus image storage and analysis software (Perkin Elmer).

### Quantitation of chromatin bridge formation

Cells were seeded in 6-well plates with glass coverslips at a seeding density of 150,000 – 200,000 cells per well, 24 h prior to the TAK-981 treatment. Media containing a range of TAK-981 concentrations was added to the cells and incubated for 24 h. Cells were fixed with 4% (w/v) formaldehyde/PBS (Pierce, 16% formaldehyde (w/v), methanol-free, Thermo Fisher, cat # 28908, diluted in PBS) for 15 min, blocked with PBG blocking buffer (0.2% fish gelatin and 0.5% (w/v) BSA, diluted in PBS) for 30 min, and incubated in blocking buffer containing 0.8 μg/ml DAPI (4’,6-diamidino-2-phenylindole, Sigma, cat # D9542) for 1 h. Cells were then mounted on glass slides using ProLong Gold Antifade mountant (Thermo Fisher, cat # P36930) and imaged on a Zeiss LSM780 confocal microscope. For quantitation of chromatin bridges, 300 – 500 nuclei per condition were observed and categorized according to presence and length of chromatin bridges, and used to calculate the proportions of nuclei displaying short or long chromatin bridges.

### Immunoprecipitation

293T cells were seeded in 10-cm plates and transfected with the plasmids expressing the NFATC2IP or NSMCE2 protein constructs that were cloned into pcDNA5-FRT/TO-3xFLAG or pcDNA3.1-C-2xHA parental vectors, respectively, by using Mirus TransIT-LT1 transfection reagent (Mirus Bio LLC, cat # MIR 2305). 24 h post-transfection, the transfection mix was removed, and cells were allowed to recover in fresh media for another 24 h before harvesting. For immunoprecipitations of FLAG-tagged peptides, whole cell lysates were prepared with high salt lysis buffer (50 mM HEPES pH 8.0, 300 nM NaCl, 2 mM EDTA pH 8.0, 0.1% (v/v) NP-40, 10% glycerol, 1x protease inhibitor cocktail (cOmplete mini, EDTA-free, Roche, cat # 11836170001) and subjected to binding of FLAG-tagged peptide species with anti-FLAG M2 magnetic beads (Sigma, cat # M8823). The purified FLAG-tagged peptides were harvested by eluting from the anti-FLAG M2 magnetic beads with 100 ng/µl 3xFLAG peptides.

### Purification of 6xHis-tagged SUMO-conjugated peptides

Cells were plated in 10-cm plates and transfected with plasmids expressing His_6_-SUMO1 (Addgene plasmid # 133770) or His_6_-SUMO2 (Addgene plasmid # 133771). 24 h post-transfection, the transfection mix was removed, and the cells were allowed to recover in fresh media for 48 h before harvest. Purifications of peptides conjugated with His_6_-SUMO1 or His_6_-SUMO2 were performed by following the protocol described in (Tatham et al. 2009), using nickel-nitrilotriacetic acid (Ni-NTA) agarose beads (QIAGEN, cat # 30230) and the manufacturer’s protocol. Briefly, cells were lysed in cell lysis buffer (6M guanidinium-HCl, 10 mM Tris-HCl, 100 mM sodium phosphate buffer pH 8.0, 20 mM NEM, 10 mM imidazole, 5 mM β-mercaptoethanol). Cell lysates were sonicated for 45 sec and cleared by centrifuging at 3000x g for 15 min at room temperature. The supernatant samples were subjected to purification of His-tagged peptides with pre-washed Ni-NTA agarose beads overnight at 4°C. Samples were then washed sequentially with (i) cell lysis buffer supplemented with 0.01% (v/v) Triton X-100, (ii) pH 8.0 wash buffer (8M urea, 10 mM Tris-HCl, 100 mM sodium phosphate buffer pH 8.0, 20 mM NEM, 10 mM imidazole, 0.1% (v/v) Triton X-100, 5 mM β-mercaptoethanol) and (iii) pH 6.3 wash buffer (8M urea, 10 mM Tris-HCl, 100 mM sodium phosphate buffer pH 6.30, 20 mM NEM, 10 mM imidazole, 0.1% (v/v) Triton X-100, 5 mM β-mercaptoethanol). Purified samples were eluted in elution buffer (200 mM imidazole, 5% (w/v) sodium dodecyl sulfate, 150 mM Tris-HCl, 30% (v/v) glycerol, 720 mM β-mercaptoethanol, 0.0025% (w/v) bromophenol blue) for 30 min at room temperature.

### Subcellular fractionation

Cells were plated in 15-cm plates, and the harvested cells were subjected to subcellular fractionations as described previously (Fradet-Turcotte et al. 2013). Briefly, cells were lysed in EBC1 buffer (50 mM Tris-HCl pH 7.50, 100 mM NaCl, 0.5% NP-40, 1 mM EDTA, 1 mM DTT, 1x protease inhibitor cocktail (cOmplete mini, EDTA-free, Roche, cat # 11836170001), 5 Mm N-ethylmaleimide (NEM, Sigma, cat # E1271)). The nuclear fraction (pellet) was separated from the cytoplasmic fraction (supernatant) by centrifuging at 1000x g for 10 min at 4°C. The nucleoplasmic fraction was obtained by resuspending the nuclear pellet in EBC2 buffer (50 mM Tris-HCl pH 7.50, 300 mM NaCl, 5 mM CaCl_2_, 1x protease inhibitor cocktail (cOmplete mini, EDTA-free, Roche, cat # 11836170001), 5 mM NEM) for 30 min on ice with occasional vortexing, after which the soluble nucleoplasmic fraction (supernatant) was separated from the insoluble chromatin (pellet) by centrifuging at 21000x g for 10 min at 4°C. The insoluble chromatin pellets were then solubilized in EBC2 buffer supplemented with micrococcal nuclease (Sigma, cat # N3755) by digesting for 45 min at 30°C. The soluble chromatin fraction samples were harvested by centrifuging at 21000x g for 10 min at 4°C and collecting the supernatant.

### Purification of SUMO-conjugated peptides

SUMO-conjugated peptides in subcellular fractionation samples were purified with biotin SUMO-Capture Reagent (Biotin S-Cap, LifeSensors Inc., cat # SM-101) following the manufacturer’s protocol. Prior to binding of SUMO-conjugated peptides, the concentrations of NP-40 in cytoplasmic fractionation samples were adjusted to 0.2% (v/v) NP-40 with dilution buffer (100 mM Tris-HCl pH 8.0, 150 mM NaCl, 5 mM EDTA pH 8.0). Samples were incubated with 1 µM Biotin S-Cap for 2 h on ice. Following the Biotin S-Cap reagent binding reaction, the input control samples were collected, and the remaining samples were incubated with Dynabeads M-280 streptavidin magnetic beads (Invitrogen, cat # 11206D) for 2 h at 4°C on an end-over-end rotator. Following the binding reaction of the Biotin S-Cap reagent to streptavidin beads, the supernatants were collected as unbound samples. The pulldown samples bound to streptavidin beads were washed 4 times with wash buffer (100 mM Tris-HCl pH 8.0, 150 mM NaCl, 5 mM EDTA pH 8.0, 0.05% (v/v) NP-40, 0.1% (v/v) Tween-20, 5 mM NEM, 1x protease inhibitor cocktail (cOmplete mini, EDTA-free, Roche, cat # 11836170001) and eluted by boiling at 95°C for 5 min.

## References

Adams J. 2001. Proteasome inhibition in cancer: development of PS-341. Semin Oncol 28: 613–619.

Adamus M, Lelkes E, Potesil D, Ganji SR, Kolesar P, Zabrady K, Zdrahal Z, Palecek JJ. 2020. Molecular Insights into the Architecture of the Human SMC5/6 Complex. J Mol Biol 432: 3820–3837.

Anwar MU, Sergeeva OA, Abrami L, Mesquita FS, Lukonin I, Amen T, Chuat A, Capolupo L, Liberali P, D’Angelo G et al. 2022. ER-Golgi-localized proteins TMED2 and TMED10 control the formation of plasma membrane lipid nanodomains. Dev Cell 57: 2334–2346 e2338.

Aragón L. 2018. The Smc5/6 Complex: New and Old Functions of the Enigmatic Long-Distance Relative. Annu Rev Genet 52: 89–107.

Becker JR, Clifford G, Bonnet C, Groth A, Wilson MD, Chapman JR. 2021. BARD1 reads H2A lysine 15 ubiquitination to direct homologous recombination. Nature.

Boddy MN, Shanahan P, McDonald WH, Lopez-Girona A, Noguchi E, Yates IJ, Russell P. 2003. Replication checkpoint kinase Cds1 regulates recombinational repair protein Rad60. Mol Cell Biol 23: 5939–5946.

Bonnon C, Wendeler MW, Paccaud JP, Hauri HP. 2010. Selective export of human GPI-anchored proteins from the endoplasmic reticulum. Journal of cell science 123: 1705–1715.

Branzei D, Sollier J, Liberi G, Zhao X, Maeda D, Seki M, Enomoto T, Ohta K, Foiani M. 2006. Ubc9- and mms21-mediated sumoylation counteracts recombinogenic events at damaged replication forks. Cell 127: 509–522.

Brinkman EK, Chen T, Amendola M, van Steensel B. 2014. Easy quantitative assessment of genome editing by sequence trace decomposition. Nucleic Acids Res 42: e168.

Cappadocia L, Lima CD. 2018. Ubiquitin-like Protein Conjugation: Structures, Chemistry, and Mechanism. Chem Rev 118: 889–918.

Chang YC, Oram MK, Bielinsky AK. 2021. SUMO-Targeted Ubiquitin Ligases and Their Functions in Maintaining Genome Stability. Int J Mol Sci 22.

Choi K, Szakal B, Chen YH, Branzei D, Zhao X. 2010. The Smc5/6 complex and Esc2 influence multiple replication-associated recombination processes in Saccharomyces cerevisiae. Mol Biol Cell 21: 2306–2314.

Colic M, Wang G, Zimmermann M, Mascall K, McLaughlin M, Bertolet L, Lenoir WF, Moffat J, Angers S, Durocher D et al. 2019. Identifying chemogenetic interactions from CRISPR screens with drugZ. Genome Med 11: 52.

Demo SD, Kirk CJ, Aujay MA, Buchholz TJ, Dajee M, Ho MN, Jiang J, Laidig GJ, Lewis ER, Parlati F et al. 2007. Antitumor activity of PR-171, a novel irreversible inhibitor of the proteasome. Cancer Res 67: 6383–6391.

Dempster JM, Rossen J, Kazachkova M, Pan J, Kugener G, Root DE, Tsherniak A. 2019. Extracting Biological Insights from the Project Achilles Genome-Scale CRISPR Screens in Cancer Cell Lines. bioRxiv: 720243.

Dewar JM, Walter JC. 2017. Mechanisms of DNA replication termination. Nat Rev Mol Cell Biol 18: 507–516.

Di Minin G, Holzner M, Grison A, Dumeau CE, Chan W, Monfort A, Jerome-Majewska LA, Roelink H, Wutz A. 2022. TMED2 binding restricts SMO to the ER and Golgi compartments. PLoS Biol 20: e3001596.

Dikic I, Schulman BA. 2022. An expanded lexicon for the ubiquitin code. Nat Rev Mol Cell Biol: 1–15.

Fenech M, Kirsch-Volders M, Natarajan AT, Surralles J, Crott JW, Parry J, Norppa H, Eastmond DA, Tucker JD, Thomas P. 2011. Molecular mechanisms of micronucleus, nucleoplasmic bridge and nuclear bud formation in mammalian and human cells. Mutagenesis 26: 125–132.

Finley D. 2009. Recognition and processing of ubiquitin-protein conjugates by the proteasome. Annu Rev Biochem 78: 477–513.

Foot N, Henshall T, Kumar S. 2017. Ubiquitination and the Regulation of Membrane Proteins. Physiol Rev 97: 253–281.

Fradet-Turcotte A, Canny MD, Escribano-Diaz C, Orthwein A, Leung CC, Huang H, Landry MC, Kitevski-Leblanc J, Noordermeer SM, Sicheri F et al. 2013. 53BP1 is a reader of the DNA-damage-induced H2A Lys 15 ubiquitin mark. Nature 499: 50–54.

Gallego-Paez LM, Tanaka H, Bando M, Takahashi M, Nozaki N, Nakato R, Shirahige K, Hirota T. 2014. Smc5/6-mediated regulation of replication progression contributes to chromosome assembly during mitosis in human cells. Mol Biol Cell 25: 302–317.

Gatti M, Pinato S, Maspero E, Soffientini P, Polo S, Penengo L. 2012. A novel ubiquitin mark at the N-terminal tail of histone H2As targeted by RNF168 ubiquitin ligase. Cell Cycle 11: 2538–2544.

Ge SX, Jung D, Yao R. 2020. ShinyGO: a graphical gene-set enrichment tool for animals and plants. Bioinformatics 36: 2628–2629.

Harper JW, Schulman BA. 2021. Cullin-RING Ubiquitin Ligase Regulatory Circuits: A Quarter Century Beyond the F-Box Hypothesis. Annu Rev Biochem 90: 403–429.

Hart T, Tong AHY, Chan K, Van Leeuwen J, Seetharaman A, Aregger M, Chandrashekhar M, Hustedt N, Seth S, Noonan A et al. 2017. Evaluation and Design of Genome-Wide CRISPR/SpCas9 Knockout Screens. G3 (Bethesda) 7: 2719–2727.

Heideker J, Prudden J, Perry JJ, Tainer JA, Boddy MN. 2011. SUMO-targeted ubiquitin ligase, Rad60, and Nse2 SUMO ligase suppress spontaneous Top1-mediated DNA damage and genome instability. PLoS Genet 7: e1001320.

Hershko A, Ciechanover A. 1998. The ubiquitin system. Annu Rev Biochem 67: 425–479.

Hodge MR, Chun HJ, Rengarajan J, Alt A, Lieberson R, Glimcher LH. 1996. NF-AT-Driven interleukin-4 transcription potentiated by NIP45. Science 274: 1903–1905.

Hoeller D, Dikic I. 2009. Targeting the ubiquitin system in cancer therapy. Nature 458: 438–444.

Hong Y, Zhang H, Gartner A. 2021. The Last Chance Saloon. Front Cell Dev Biol 9: 671297.

Hou W, Gupta S, Beauchamp MC, Yuan L, Jerome-Majewska LA. 2017. Non-alcoholic fatty liver disease in mice with heterozygous mutation in TMED2. PLoS One 12: e0182995.

Hyer ML, Milhollen MA, Ciavarri J, Fleming P, Traore T, Sappal D, Huck J, Shi J, Gavin J, Brownell J et al. 2018. A small-molecule inhibitor of the ubiquitin activating enzyme for cancer treatment. Nat Med 24: 186–193.

Jackson SP, Durocher D. 2013. Regulation of DNA damage responses by ubiquitin and SUMO. Mol Cell 49: 795–807.

Jacome A, Gutierrez-Martinez P, Schiavoni F, Tenaglia E, Martinez P, Rodriguez-Acebes S, Lecona E, Murga M, Mendez J, Blasco MA et al. 2015. NSMCE2 suppresses cancer and aging in mice independently of its SUMO ligase activity. EMBO J 34: 2604–2619.

Joazeiro CAP. 2017. Ribosomal Stalling During Translation: Providing Substrates for Ribosome-Associated Protein Quality Control. Annu Rev Cell Dev Biol 33: 343–368.

Langston SP, Grossman S, England D, Afroze R, Bence N, Bowman D, Bump N, Chau R, Chuang BC, Claiborne C et al. 2021. Discovery of TAK-981, a First-in-Class Inhibitor of SUMO-Activating Enzyme for the Treatment of Cancer. J Med Chem 64: 2501–2520.

Li S, Bonner JN, Wan B, So S, Mutchler A, Gonzalez L, Xue X, Zhao X. 2021. Esc2 orchestrates substrate-specific sumoylation by acting as a SUMO E2 cofactor in genome maintenance. Genes Dev 35: 261–272.

Lightcap ES, Yu P, Grossman S, Song K, Khattar M, Xega K, He X, Gavin JM, Imaichi H, Garnsey JJ et al. 2021. A small-molecule SUMOylation inhibitor activates antitumor immune responses and potentiates immune therapies in preclinical models. Sci Transl Med 13: eaba7791.

Mattiroli F, Vissers JH, van Dijk WJ, Ikpa P, Citterio E, Vermeulen W, Marteijn JA, Sixma TK. 2012. RNF168 Ubiquitinates K13-15 on H2A/H2AX to Drive DNA Damage Signaling. Cell 150: 1182–1195.

Matunis MJ, Coutavas E, Blobel G. 1996. A novel ubiquitin-like modification modulates the partitioning of the Ran-GTPase-activating protein RanGAP1 between the cytosol and the nuclear pore complex. J Cell Biol 135: 1457–1470.

Meyer H, Weihl CC. 2014. The VCP/p97 system at a glance: connecting cellular function to disease pathogenesis. Journal of cell science 127: 3877–3883.

Miyabe I, Morishita T, Hishida T, Yonei S, Shinagawa H. 2006. Rhp51-dependent recombination intermediates that do not generate checkpoint signal are accumulated in Schizosaccharomyces pombe rad60 and smc5/6 mutants after release from replication arrest. Mol Cell Biol 26: 343–353.

Morishita T, Tsutsui Y, Iwasaki H, Shinagawa H. 2002. The Schizosaccharomyces pombe rad60 gene is essential for repairing double-strand DNA breaks spontaneously occurring during replication and induced by DNA-damaging agents. Mol Cell Biol 22: 3537–3548.

Northrop A, Byers HA, Radhakrishnan SK. 2020. Regulation of NRF1, a master transcription factor of proteasome genes: implications for cancer and neurodegeneration. Mol Biol Cell 31: 2158–2163.

Novatchkova M, Bachmair A, Eisenhaber B, Eisenhaber F. 2005. Proteins with two SUMO-like domains in chromatin-associated complexes: the RENi (Rad60-Esc2-NIP45) family. BMC Bioinformatics 6: 22.

Olivieri M, Cho T, Alvarez-Quilon A, Li K, Schellenberg MJ, Zimmermann M, Hustedt N, Rossi SE, Adam S, Melo H et al. 2020. A Genetic Map of the Response to DNA Damage in Human Cells. Cell 182: 481–496 e421.

Olivieri M, Durocher D. 2021. Genome-scale chemogenomic CRISPR screens in human cells using the TKOv3 library. STAR Protoc 2: 100321.

Oughtred R, Rust J, Chang C, Breitkreutz BJ, Stark C, Willems A, Boucher L, Leung G, Kolas N, Zhang F et al. 2021. The BioGRID database: A comprehensive biomedical resource of curated protein, genetic, and chemical interactions. Protein Sci 30: 187–200.

Park JM, Yang SW, Yu KR, Ka SH, Lee SW, Seol JH, Jeon YJ, Chung CH. 2014. Modification of PCNA by ISG15 plays a crucial role in termination of error-prone translesion DNA synthesis. Mol Cell 54: 626–638.

Payne F, Colnaghi R, Rocha N, Seth A, Harris J, Carpenter G, Bottomley WE, Wheeler E, Wong S, Saudek V et al. 2014. Hypomorphism in human NSMCE2 linked to primordial dwarfism and insulin resistance. J Clin Invest 124: 4028–4038.

Potts PR, Yu H. 2005. Human MMS21/NSE2 is a SUMO ligase required for DNA repair. Mol Cell Biol 25: 7021–7032.

Prudden J, Perry JJ, Arvai AS, Tainer JA, Boddy MN. 2009. Molecular mimicry of SUMO promotes DNA repair. Nat Struct Mol Biol 16: 509–516.

Prudden J, Perry JJ, Nie M, Vashisht AA, Arvai AS, Hitomi C, Guenther G, Wohlschlegel JA, Tainer JA, Boddy MN. 2011. DNA repair and global sumoylation are regulated by distinct Ubc9 noncovalent complexes. Mol Cell Biol 31: 2299–2310.

Pryzhkova MV, Jordan PW. 2016. Conditional mutation of Smc5 in mouse embryonic stem cells perturbs condensin localization and mitotic progression. Journal of cell science 129: 1619–1634.

Psakhye I, Jentsch S. 2012. Protein Group Modification and Synergy in the SUMO Pathway as Exemplified in DNA Repair. Cell 151: 807–820.

Radhakrishnan SK, den Besten W, Deshaies RJ. 2014. p97-dependent retrotranslocation and proteolytic processing govern formation of active Nrf1 upon proteasome inhibition. Elife 3: e01856.

Raschle M, Smeenk G, Hansen RK, Temu T, Oka Y, Hein MY, Nagaraj N, Long DT, Walter JC, Hofmann K et al. 2015. DNA repair. Proteomics reveals dynamic assembly of repair complexes during bypass of DNA cross-links. Science 348: 1253671.

Raso MC, Djoric N, Walser F, Hess S, Schmid FM, Burger S, Knobeloch KP, Penengo L. 2020. Interferon-stimulated gene 15 accelerates replication fork progression inducing chromosomal breakage. J Cell Biol 219.

Robey RW, Pluchino KM, Hall MD, Fojo AT, Bates SE, Gottesman MM. 2018. Revisiting the role of ABC transporters in multidrug-resistant cancer. Nat Rev Cancer 18: 452–464.

Ruvkun G, Lehrbach N. 2023. Regulation and Functions of the ER-Associated Nrf1 Transcription Factor. Cold Spring Harb Perspect Biol 15.

Sebesta M, Urulangodi M, Stefanovie B, Szakal B, Pacesa M, Lisby M, Branzei D, Krejci L. 2017. Esc2 promotes Mus81 complex-activity via its SUMO-like and DNA binding domains. Nucleic Acids Res 45: 215–230.

Sekiyama N, Arita K, Ikeda Y, Hashiguchi K, Ariyoshi M, Tochio H, Saitoh H, Shirakawa M. 2010. Structural basis for regulation of poly-SUMO chain by a SUMO-like domain of Nip45. Proteins 78: 1491–1502.

Sifri C, Hoeg L, Durocher D, Setiaputra D. 2023. An AlphaFold2 map of the 53BP1 pathway identifies a direct SHLD3-RIF1 interaction critical for DNA repair activity. bioRxiv: 2023.2001.2012.523815.

Sollier J, Driscoll R, Castellucci F, Foiani M, Jackson SP, Branzei D. 2009. The Saccharomyces cerevisiae Esc2 and Smc5-6 proteins promote sister chromatid junction-mediated intra-S repair. Mol Biol Cell 20: 1671–1682.

Soucy TA, Smith PG, Milhollen MA, Berger AJ, Gavin JM, Adhikari S, Brownell JE, Burke KE, Cardin DP, Critchley S et al. 2009. An inhibitor of NEDD8-activating enzyme as a new approach to treat cancer. Nature 458: 732–736.

Steffen J, Seeger M, Koch A, Kruger E. 2010. Proteasomal degradation is transcriptionally controlled by TCF11 via an ERAD-dependent feedback loop. Mol Cell 40: 147–158.

Szklarczyk D, Gable AL, Nastou KC, Lyon D, Kirsch R, Pyysalo S, Doncheva NT, Legeay M, Fang T, Bork P et al. 2021. The STRING database in 2021: customizable protein-protein networks, and functional characterization of user-uploaded gene/measurement sets. Nucleic Acids Res 49: D605–D612.

Tatham MH, Geoffroy MC, Shen L, Plechanovova A, Hattersley N, Jaffray EG, Palvimo JJ, Hay RT. 2008. RNF4 is a poly-SUMO-specific E3 ubiquitin ligase required for arsenic-induced PML degradation. Nat Cell Biol 10: 538–546.

Tatham MH, Rodriguez MS, Xirodimas DP, Hay RT. 2009. Detection of protein SUMOylation in vivo. Nat Protoc 4: 1363–1371.

Tsitsiridis G, Steinkamp R, Giurgiu M, Brauner B, Fobo G, Frishman G, Montrone C, Ruepp A. 2023. CORUM: the comprehensive resource of mammalian protein complexes-2022. Nucleic Acids Res 51: D539–D545.

Ulrich HD, Vogel S, Davies AA. 2005. SUMO keeps a check on recombination during DNA replication. Cell Cycle 4: 1699–1702.

Urulangodi M, Sebesta M, Menolfi D, Szakal B, Sollier J, Sisakova A, Krejci L, Branzei D. 2015. Local regulation of the Srs2 helicase by the SUMO-like domain protein Esc2 promotes recombination at sites of stalled replication. Genes Dev 29: 2067–2080.

Varejao N, Lascorz J, Codina-Fabra J, Belli G, Borras-Gas H, Torres-Rosell J, Reverter D. 2021. Structural basis for the E3 ligase activity enhancement of yeast Nse2 by SUMO-interacting motifs. Nat Commun 12: 7013.

Venegas AB, Natsume T, Kanemaki M, Hickson ID. 2020. Inducible Degradation of the Human SMC5/6 Complex Reveals an Essential Role Only during Interphase. Cell Reports 31: 107533.

Vertegaal ACO. 2022. Signalling mechanisms and cellular functions of SUMO. Nat Rev Mol Cell Biol 23: 715–731.

Vondrova L, Kolesar P, Adamus M, Nociar M, Oliver AW, Palecek JJ. 2020. A role of the Nse4 kleisin and Nse1/Nse3 KITE subunits in the ATPase cycle of SMC5/6. Sci Rep 10: 9694.

Wardlaw CP, Petrini JHJ. 2022. ISG15 conjugation to proteins on nascent DNA mitigates DNA replication stress. Nat Commun 13: 5971.

Warecki B, Sullivan W. 2020. Mechanisms driving acentric chromosome transmission. Chromosome Res 28: 229–246.

Wertz IE, Wang X. 2019. From Discovery to Bedside: Targeting the Ubiquitin System. Cell Chem Biol 26: 156–177.

Wilson MD, Benlekbir S, Fradet-Turcotte A, Sherker A, Julien JP, McEwan A, Noordermeer SM, Sicheri F, Rubinstein JL, Durocher D. 2016. The structural basis of modified nucleosome recognition by 53BP1. Nature 536: 100–103.

Ye Y, Tang WK, Zhang T, Xia D. 2017. A Mighty “Protein Extractor” of the Cell: Structure and Function of the p97/CDC48 ATPase. Front Mol Biosci 4: 39.

Yu Y, Li S, Ser Z, Kuang H, Than T, Guan D, Zhao X, Patel DJ. 2022. Cryo-EM structure of DNA-bound Smc5/6 reveals DNA clamping enabled by multi-subunit conformational changes. Proc Natl Acad Sci U S A 119: e2202799119.

Yu Y, Li S, Ser Z, Sanyal T, Choi K, Wan B, Kuang H, Sali A, Kentsis A, Patel DJ et al. 2021. Integrative analysis reveals unique structural and functional features of the Smc5/6 complex. Proc Natl Acad Sci U S A 118.

Zhang H, Jin X, Huang H. 2023. Deregulation of SPOP in Cancer. Cancer Res 83: 489–499.

Zheng N, Shabek N. 2017. Ubiquitin Ligases: Structure, Function, and Regulation. Annu Rev Biochem 86: 129–157.

Zhou HJ, Wang J, Yao B, Wong S, Djakovic S, Kumar B, Rice J, Valle E, Soriano F, Menon MK et al. 2015. Discovery of a First-in-Class, Potent, Selective, and Orally Bioavailable Inhibitor of the p97 AAA ATPase (CB-5083). J Med Chem 58: 9480–9497.

Zimmermann M, Murina O, Reijns MAM, Agathanggelou A, Challis R, Tarnauskaite Z, Muir M, Fluteau A, Aregger M, McEwan A et al. 2018. CRISPR screens identify genomic ribonucleotides as a source of PARP-trapping lesions. Nature 559: 285–289.

